# Ahead of the membrane curve: *in silico* insights into amyloid-*β* aggregation

**DOI:** 10.64898/2026.08.22.746319

**Authors:** Pedro Maximiano, Mohtadin Hashemi

## Abstract

Membrane surfaces can accelerate amyloid *β* (A*β*) aggregation, yet the role of membrane curvature in this process remains poorly understood. Here, we used multi-million atom all-atom molecular dynamics simulations to compare the adsorption, conformational dynamics, and oligomerization of four A*β*42 peptides at a planar neuronal membrane and a highly curved lipid vesicle. For both systems, all peptides adsorbed within the first 2 *µ*s, but their subsequent behavior differed substantially. The curved membrane exhibited a larger area per lipid and more extensive hydrophobic packing defects, allowing A*β*42 to penetrate more deeply and form strong contacts with lipid tails through its central hydrophobic core and C-terminal region. These interactions disrupted a solution-formed dimer and limited peptide-peptide association during the simulated interval. Additionally, vesicle-bound peptides adopted more extended conformations with increased *β*-structure and *β*-hairpin formation compared with peptides at the planar membrane. A*β*42 adsorption was also corelated to lipid reorganization in the vesicle. In contrast, the planar membrane supported weaker adsorption and stable dimer-to-trimer growth but showed little large-scale lipid segregation. These findings reveal that curvature reshapes the early A*β*42 aggregation landscape by strengthening peptide-lipid interactions, altering aggregation-prone conformations, and reorganizing membrane domains. Membrane geometry should therefore be considered alongside lipid composition in mechanistic models of A*β*42 oligomerization and membrane-associated toxicity.

## Introduction

Alzheimer’s disease (AD), the leading cause of dementia, is an escalating global health and socioeconomic challenge. The World Health Organization (WHO) estimates that 57 million people lived with dementia in 2021, with nearly 10 million new cases each year ^1^. The prevalence of AD is projected to increase to approximately 152.8 million by 2050^2^, imposing a significant socio-economic burden^3^.

The pathophysiology of AD is marked by the extracellular accumulation of amyloid-*β* (A*β*) peptides into plaques and intracellular tau neurofibrillary tangles, leading to synaptic disruption and neuronal loss^4–6^. The A*β* peptide is produced by the sequential proteolytic cleavage of amyloid precursor protein (APP) by *β*-and *γ*-secretases^6^. While A*β*40 is the most abundant isoform, A*β*42, which contains two additional hydrophobic residues at the C-terminus, is significantly more aggregation-prone and neurotoxic^7,8^.

Although mature amyloid fibrils are a hallmark of AD, converging evidence identifies small, soluble A*β* oligomers as highly neurotoxic species that drive synaptic dysfunction and early cognitive decline, placing the initial steps of aggregation at the center of mechanistic and therapeutic interest^9–11^. A central challenge in the field is that A*β* oligomers are transient, heterogeneous, and sensitive to the chemical milieu.

Soluble A*β*42 monomers in the extracellular space first adsorb onto the neuronal membrane surface. Once bound, their diffusion is confined to the plane of the bilayer, which increases the local concentration of the monomers, thus enhancing the probability of their interaction to form a stable nucleus^12^. Recent works provide strong evidence that the surface of the cell membrane catalyzes the formation of A*β* oligomers^13–17^. Experimental studies using supported lipid bilayers have shown that membrane contact can accelerate A*β* aggregation, even at physiologically relevant low nanomolar concentrations, consistent with surface-mediated pathways in which local enrichment and interfacial templating reduce kinetic barriers^13,18–20^. Transient interactions with the lipid interface drive A*β* monomer misfolding by stabilizing aggregation-prone conformations featuring *β* secondary structures^13,19^. Once formed, surface-assembled oligomers do not remain permanently anchored to the bilayer; they are capable of spontaneously dissociating back into the aqueous environment, acting as mobile, neurotoxic seeds that further propagate the amyloid cascade^13^. It was shown that the aggregation process at the surface depends heavily on lipid composition, being particularly sensitive to the presence of cholesterol and gangliosides in the membrane^13,21–24^.

Besides composition, curvature is understood as a critical parameter governing the biological functions of the cell membrane. This is particularly true of neuronal compartments, as changes in curvature are key to synaptic physiology, in which highly curved dendritic spines and synaptic vesicles are abundant. Because the flat membrane models traditionally used in computational and *in vitro* studies fail to capture the topographical complexity of a living neuron, understanding how high-curvature environments dictate protein-lipid interactions is critical. Curvature can alter protein binding, insertion, and peptide conformational equilibria, which can in turn promote lipid reorganization and domain behavior, feeding back into oligomerization pathways^25^.

Experimental studies show that these curved interfaces may directly stabilize aggregation-prone conformations by inducing *β*-structure in A*β* monomers^26,27^. Rates of fibrillation A*β* were found to increase in the presence of vesicles of increasing curvature (decreasing radius)^27–29^. Specifically, high-curvature vesicles possess distinct lipid packing defects that expose hydrophobic clefts, which offer binding sites that can effectively concentrate A*β* monomers and promote productive primary nucleation, compared to low-curvature bilayers^28–30^. Furthermore, in the presence of highly curved lipid vesicles, the catalytic effect extends beyond initial oligomerization; the curved lipid surface actively interacts with preformed aggregates to reinforce secondary nucleation pathways, thereby accelerating overall fibril formation^18^. Despite these isolated findings, the field lacks a systematic, high-resolution framework linking membrane curvature to the formation, stability, and membrane activity of early A*β* oligomers.

To address this gap, we employed microsecond-scale, all-atom molecular dynamics (MD) simulations to systematically investigate the aggregation dynamics of A*β*42 monomers in the presence of distinct membrane geometries. By comparing a flat bilayer system with a highly curved vesicle model, we aim to elucidate how membrane curvature dictates A*β*42 adsorption, conformational conversion, and the mechanism of surface-catalyzed aggregation. Furthermore, we investigated the reciprocal effect of A*β*42 binding on the organization of the lipid bilayer, highlighting the coupled dynamics of protein misfolding and curvature-induced lipid raft formation.

## Methods

### Model construction and simulation details

Two lipid bilayer models with distinct curvatures – a double flat bilayer and a vesicle (spherical bilayer) – were constructed, both in the absence and presence of A*β*42. The composition of the simulation systems was chosen to emulate physiological conditions as closely as possible.

The lipid compositions followed a simplified neuronal plasma membrane model proposed by Wilson et al.^25^, which reproduces the dynamics of the more complex membrane described by Ingólfsson et al. ^31^. The inner leaflet consisted of 44.7% cholesterol, 15.9% 1-palmitoyl-2-oleoyl-sn-glycero-3-phosphocholine (POPC), 25.1% 1-palmitoyl-2-docosahexaenoyl-sn-glycero-3-phosphoethanolamine (PUPE), 11.7% 1-palmitoyl-2-arachidonoyl-sn-glycero-3-phospho-L-serine (PAPS), and 2.7% N-stearoyl-D-erythrosphingosylphosphorylcholine (DPSM). The outer leaflet contained 44.6% cholesterol, 26.2% POPC, 11.7% PUPE, 9.5% DPSM, and 8.0% N-stearoyl-*β*-D-galactosylceramide (DPGS) (lipid structures are schematized in Figure SI.1).

The flat bilayer model was constructed in the CHARMM-GUI platform^32,33^ and two copies of it were placed along the *z* axis of a 33.3 × 33.3 × 33.0 nm box, with a 10 nm spacing between them. On the other hand, a vesicle of diameter ∼21 nm was built with the CELLMicrocosmos 2.2 MembraneEditor^34^ tool (with leaflet lipid counts estimated from CHARMM-GUI^35^), and placed at the center of a cubic box of 35 nm side length. This system was then solvated and neutralized with K^+^ ions, followed by minimization and relaxation under an NPT ensemble, until an equilibrated vesicle structure was obtained. During the equilibration, some cholesterol molecules migrated between leaflets, but the overall molar percentages of each lipid remained very close to the initial values (see detailed lipid bilayer compositions in Table SI.1). The solvent and ions were subsequently removed prior to assembling the full vesicle system.

The space inside the vesicle, or between the two flat bilayers, marks the intracellular domain. Glutamate neurotransmitter molecules were placed in this region at a concentration of 0.7 M^36^, in order to mimic fully loaded synaptic vesicles. The remaining free volume in the box thus constitutes the extracellular domain. Na^+^ and K^+^ ions were inserted on each domain at their corresponding physiological concentrations^37^. The total number of Na^+^ and K^+^ ions placed on the intracellular domain was chosen to neutralize the negative charge of glutamate and PAPS molecules. On the extracellular domain, charge neutrality was accomplished through the addition of Cl ^-^ counterions. Finally, each system was fully solvated. The final molecular composition of each system is summarized in Table SI.1.

Once assembled, both systems were equilibrated in the GROMACS 2024.3^38^ package, with the following protocol: (i) energy minimization with position restraints on the lipid headgroups and ions; (ii) unrestrained energy minimization; (iii) annealing to 300 K (with position restraints); (iv) equilibration in the NVT ensemble at 300 K (with position restraints); (v) equilibration in the NPT ensemble (300 K, 1 bar) with position restraints gradually removed; (vi) final unrestrained NPT equilibration at 300 K and 1 bar for 15 ns.

The equilibrated system coordinates, along with topologies, were then converted to Desmond format (dms), and submitted to a 1 µs-long MD run in the Anton3^39^ supercomputer.

The final Anton3 frames were used to construct the systems with A*β*42. To that end, four A*β*42 monomers were inserted in the extracellular domain with random orientations by replacing water molecules. Their centers-of-mass (COM) were positioned 5 nm away from the membrane surface and at least 14 nm away from one another. The structure of the A*β*42 monomers was derived from MD simulations by Hashemi et al.^40^, and includes an additional C-terminal cysteine residue commonly used for covalent anchoring in single-molecule force spectroscopy experiments. Due to the net negative charge of the A*β*42 monomers, additional Na^+^ and K^+^ ions were added (at the extracellular molar ratio) to restore neutrality (final compositions are detailed in Table SI.1). The systems were then minimized and equilibrated in GROMACS 2024.3 following the protocol described previously, with position restraints applied only to the A*β*42 backbone atoms during stages (i)–(v). Production simulations were subsequently performed for 23 µs on Anton3. Initial and final frames of the trajectories are depicted in Figure SI.2.

All simulations employed the CHARMM36m^41^ force field for proteins and CHARMM36^42^ for lipids. Parameters for glutamate molecules were drawn from CGenFF^43^, whereas the TIP3P model was used for water. A cutoff of 1.2 nm was applied to all non-bonded interactions. Long-range treatment of electrostatic interactions was accomplished in GROMACS and Anton3 through the Particle Mesh Ewald (PME)^44^ and Particle Mesh *u*-series^45^ algorithms, respectively. All Anton 3 production runs were performed in the NPT ensemble at 298.15 K and 1 bar, maintained through coupling to an antithetic thermostat^46^ and a Monte-Carlo barostat^47^, with semi-isotropic or isotropic coupling applied to flat bilayer and vesicle systems, respectively. An integration time step of 2.0 fs was used with the multigrator scheme^46^, while equilibration runs on GROMACS employed the leapfrog integrator with the same time step.

### Trajectory analysis

The geometric properties of the lipid membranes, namely the area per lipid (APL) and membrane thickness, were determined following leaflet assignment. Lipids were first classified as belonging to the inner or outer leaflet by computing the positions of representative headgroup atoms (P for phospholipids, C1 for the *β*-galactose headgroup of DPGS, and the O atom for cholesterol) along the direction normal to the membrane (*z* axis for the double flat bilayer and radial coordinate for the vesicle). A two-component Gaussian mixture model (GMM) was then fitted to the distribution of these positions. Each leaflet was associated with one of the Gaussian components, and a midplane threshold was defined at the minimum of the GMM between the two peaks. To improve robustness, outliers in this scheme, which led to incorrect leaflet membership, were reassigned using a neighbor-based correction algorithm.

Following leaflet assignment, the membrane thickness was computed as the distance between the average headgroup positions of the two leaflets. The APL was calculated for each lipid via a Voronoi tessellation projected onto the leaflet surface (planar for bilayers and spherical for vesicles). Lipid acyl chain order parameters were computed using the gorder tool^48^ according to Equation 1:

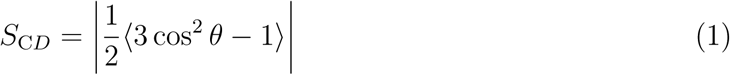

where *θ* is the angle between the acyl C–H bond vector and the local membrane normal, evaluated at each frame.

Lipid hydrophobic packing defects were computed via the procedure outlined by Cui et al.^49^. Briefly, exposed areas of the bilayer hydrophobic domain were identified using a solvent-accessible surface area (SASA) algorithm, implement via the measure sasa tool in VMD^50^, with a probe radius of 3.0 Å and 10000 samples. The resulting points were clustered in two or three dimensions (neighbor cutoff of 1.8 Å), and then projected onto a cartesian grid (*xy* plane) or a hierarchical equal area isolatitude pixelation (HEALPix) spherical grid, for bilayer or vesicle systems, respectively, using a resolution of 0.16 Å^2^. Defect areas were obtained by counting the number of occupied pixels per cluster.

Protein–protein and protein–lipid contacts were defined whenever the distance between two heavy atoms was less than 4.5 Å^51^, whereas hydrophobic contacts followed the same criteria, but considering only C atoms in hydrophobic moieties. A minimum of 50 total contacts were considered as the threshold to define the formation of aggregates between two A*β*42 molecules. The donor-acceptor distance and donor-hydrogen-acceptor angle cutoffs used to identify hydrogen bonds (H-bonds) were 3.0 Å and 135°, respectively.

The secondary structure of each A*β*42 peptide was determined using the DSSP algorithm^52^. *β*-hairpin motifs were identified by detecting contiguous *β*-strand segments of at least two residues – required for stable secondary structure formation^53^ – separated by a turn of 2–5 residues, consistent with canonical geometries described by Sibanda and Thornton^54^. A minimum of 15 heavy atom contacts (cutoff of 4.5 Å) between the strands was enforced to distinguish hairpins from *β*-arch motifs^55^.

All analysis were performed with custom scripts employing routines from the CPP-TRAJ^56^ (version 6.29.13) and MDAnalysis^57,58^ (version 2.9.0) softwares. Snapshots of molecular structures and trajectory frames were generated with VMD^59,60^.

## Results and discussion

### Lipid-protein interactions

Lipid bilayer curvature was found to strongly influence the adsorption of A*β*42 peptides. Figure 1 shows the time evolution of the peptide COM positions along the direction normal to the membrane. In both systems, all peptides adsorbed within the first 2 µs. However, for the flat bilayer, almost all A*β*42 chains – except A*β*42-2 – remained above the membrane surface, whereas in the vesicle system all peptides were able to penetrate deeper beneath the outer surface. This observation is further supported by the average residue positions computed over the final 1 µs of the simulations (Figure SI.3), which show that A*β*42 peptides are partially inserted in the inner domain of the curved membrane, while mostly failing to do so in the flat geometry.

**Figure 1.**
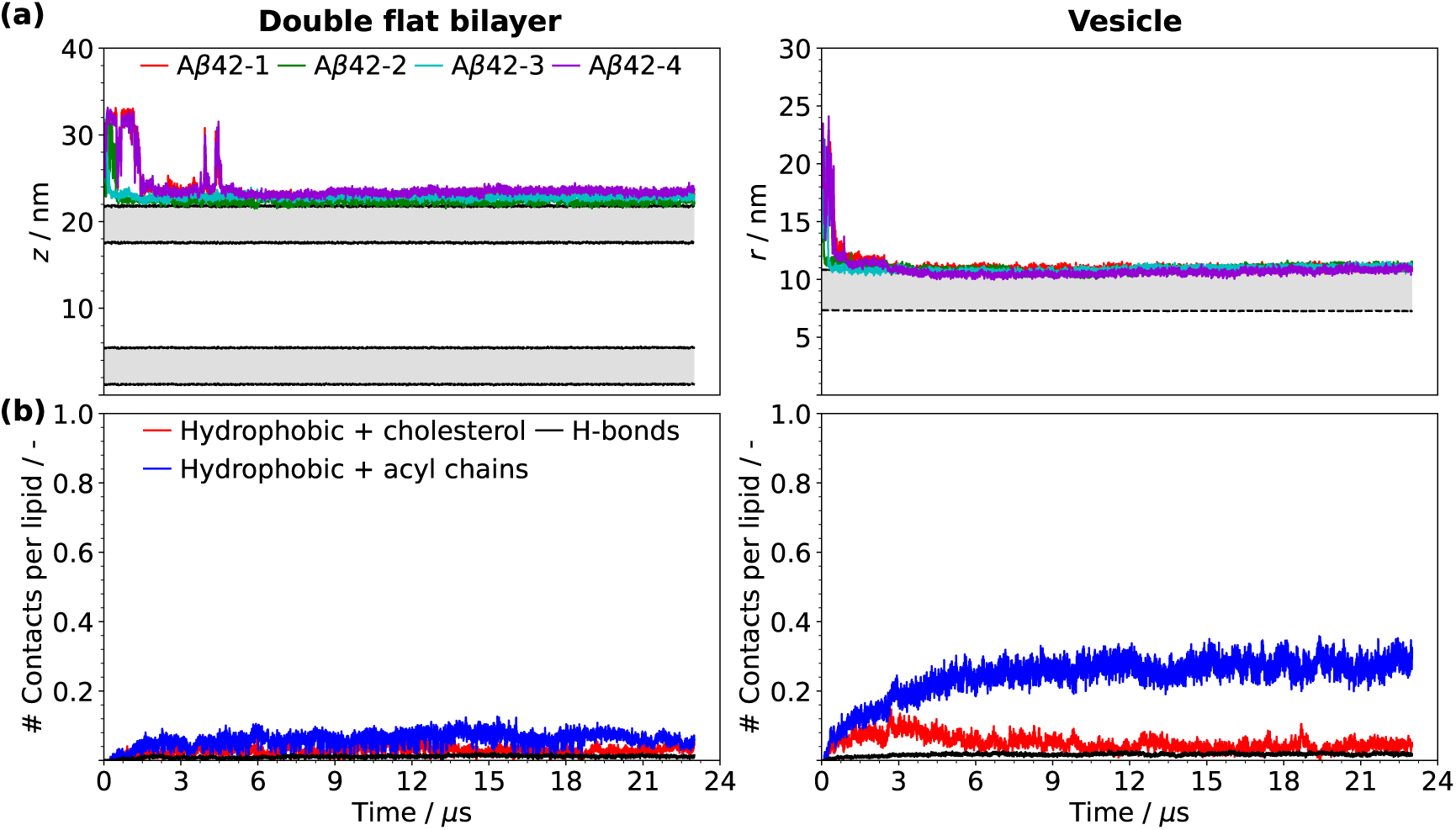
Position of COM of A*β*42 peptides in the direction transversal to the lipid bilayer(s) (a) and number of H-bonds and hydrophobic contacts between A*β*42 and lipid molecules (b) recorded for the flat bilayer (left) and vesicle (right) trajectories. In the top plots, the grey area represents the position of the membrane, with the dotted black line marking the average lipid headgroup positions. Colors of each A*β*42 peptide in (a) match those in the trajectory frames shown in Figure SI.2.

The enhanced adsorption of A*β*42 to the vesicle is primarily driven by hydrophobic interactions with lipid molecules, particularly with the acyl chains (Figure 1 (b)). In comparison, hydrophobic contacts with cholesterol and H-bonds involving lipid headgroups contribute only marginally. By contrast, interactions between A*β*42 and lipid acyl chains of the planar bilayer are significantly weaker and occur at levels comparable to hydrogen bonding.

In the vesicle system, A*β*42 interacts preferentially with lipid acyl chains through its central hydrophobic core (CHC, residues 17–21) and hydrophobic C-terminal (residues 30– 42) regions. This is evidenced by the average positions of the corresponding residues, which lie below the vesicle surface (Figure SI.3 (b)). A similar behavior is observed for the only peptide in the flat bilayer system that partially penetrated the membrane surface, A*β*42-2 (Figure SI.3 (a)). The average protein-lipid contact matrix (Figure SI.4 (b)) not only confirms the dominant role of the CHC and C-terminal domains in mediating interactions with the membrane, but also pinpoints the acyl chains of PUPE, DPSM and DPGS as the most favored partners for such interactions.

The N-terminal metal-binding region (residues 1–16) also possess sites that contribute significantly to interactions with the vesicle. In particular, Phe4 and Tyr10 form hydrophobic contacts with PUPE, DPSM, and DPGS acyl chains, and can penetrate below the vesicle surface (Figure SI.3 (b)). In addition, polar residues such as Arg5, Asp7, and Glu11 form H-bonds with the DPGS headgroups. These interactions (along with further contributions from Gln16) are also observed in the flat bilayer system (Figure SI.4 (a)), indicating that the *β*-galactose headgroup of DPGS plays an important role in anchoring A*β*42 to the membrane surface regardless of curvature. Experimental studies have similarly shown that the metal-binding domain of A*β*42 can selectively bind to the *β*-galactose-group of DGPS (Gal-Cer), with Arg5 and Tyr10 also identified as key residues in this process^61,62^. Notably, the A*β*42–*β*-galactose interactions were more pronounced in the vesicle system compared to its flat counterpart (Figure SI.4), suggesting a greater accessibility of the *β*-galactose at the surface facilitated by its curvature. After DPGS, another notable binding partner for A*β*42 is the PUPE lipid species. Not only are its acyl chains heavily engaged with the CHC and hydrophobic domains of A*β*42 via hydrophobic contacts, but electrostatic interactions arise between negatively charged Glu and Asp amino acids in the polar domain and the positively charged NH_3_^+^ moieties in the phosphoetanolamine headgroup. Finally, POPC also participates in significant hydrophobic interactions with A*β*42, albeit to a lesser extent than PUPE. This observation aligns with the results of MD simulations by Tofelanu et al. ^63^, which showed stronger binding of A*β* to 1-palmitoyl-2-oleoyl-sn-glycero-3-phosphoethanolamine (POPE) compared to POPC. The headgroups of these two lipids also provide notable interaction points for A*β*42 in a flat system, albeit the interactions are much weaker compared to the vesicle system (Figure SI.4 (a)).

The stronger hydrophobic interactions observed in the vesicle arise from its highly curved surface, which exhibits a larger APL compared to the outer leaflet of the flat bilayer, as seen in Figure 2 (a) (averages computed every 100 ns from the time series depicted in Figures SI.5 and SI.6). A higher APL reflects increased exposure of the membrane hydrophobic core, facilitating interactions with the hydrophobic regions of A*β*42. This interpretation is supported by the analysis of hydrophobic packing defects (Figures 2 (b)–(f)). Vesicle systems consistently display a higher density of defects and a larger fraction of defect area compared to flat bilayers, irrespective of peptide presence. Moreover, the defect area distributions (Figure SI.7) extend to larger values for the vesicle and feature smaller exponential decay constants, which is indicative of a surface populated by defects with larger average areas.

**Figure 2.**
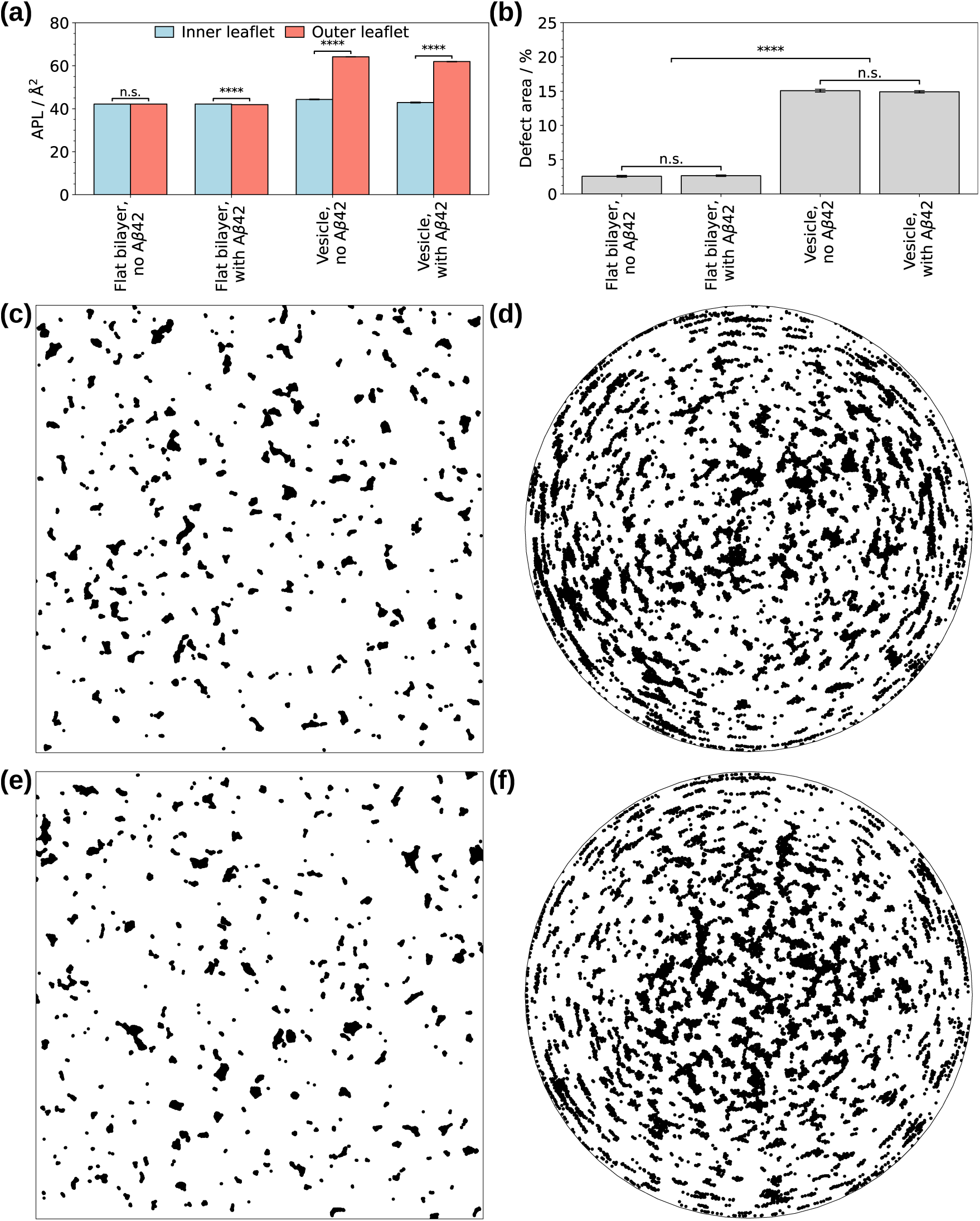
(a) Average APL of outer and inner leaflets and (b) average percentage of lipid packing defect area for the simulated systems. Distribution of lipid hydrophobic packing defect sites at the last trajectory frame for the flat bilayer systems, without (c) and with (e) A*β*42, and for the vesicle systems, without (d) and with (f) A*β*42. For (a) and (b) statistics were collected for 10 frames, taken from the end of the trajectories at every 100 ns (corresponding to the decay time of the autocorrelation function of APL). A two-sample Kolomogorov-Smirnov test was performed for assessing differences between the distributions of pairs of datasets, with **** indicating statistical significance at *p <* 0.0001 and n.s. *p >* 0.1. (c) and (e) represent normalized cartesian coordinates projected on the *xy* plane, whereas (d) and (f) represent polar coordinates in a Lambert azimuthal equal-area projection.^12^

### Effects of curvature on A***β***42 dynamics

The curvature-dependent differences in A*β*42 adsorption also have significant implications for peptide aggregation and structural evolution upon membrane binding. Figure 3 (a) tracks the oligomerization state of A*β*42 over time for both membrane geometries. In both systems, a dimer forms spontaneously in solution within the first 2 µs and subsequently migrates to-ward the membrane surface (Figure 1 (a)). In the flat bilayer system, this dimer remains stable throughout the simulation, and a trimer forms in the final 4 µs via the addition of a third monomer. By contrast, upon adsorption onto the vesicle, the dimer dissociates as strong peptide–lipid interactions develop (Figure 1 (b)). As shown in Figure SI.8, the formation of hydrophobic contacts between A*β*42 and lipid acyl chains weakens intermolecular hydrophobic interactions between peptides (rather than intramolecular contacts), which are responsible for stabilizing the dimer.

**Figure 3.**
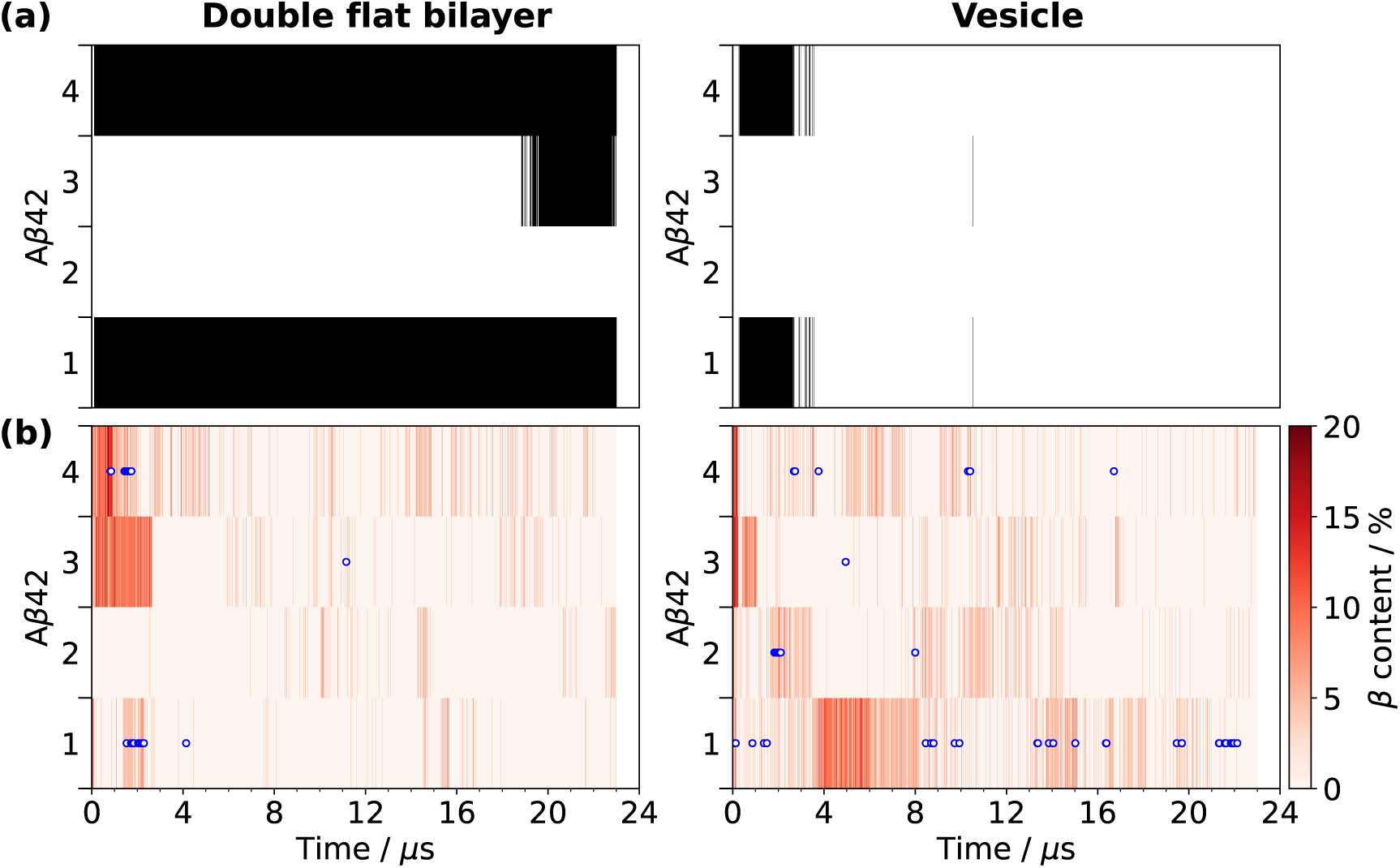
State of oligomerization (a) and percentage of *β* secondary structures (b) of each A*β*42 peptide recorded for the flat bilayer (left) and vesicle (right) trajectories. In (a), black pixels represent A*β*42 peptides involved in the same oligomer. In (b), blue and white circles represent frames for which *β*-hairpin motifs were identified.

The emergence of *β*-secondary structures is a key step in A*β*42 misfolding and sub-sequent oligomer and fibril formation ^64–67^. The time evolution of the *β*-structure content (including *β*-sheets and *β*-bridges) is shown in Figure 3 (b), with full assignments provided in Figure SI.9. Prior to adsorption, peptides in both systems display comparable *β*-content.

However, after adsorption, a clear divergence emerges, as A*β*42 monomers bound to the vesicle exhibit a higher probability of adopting *β*-structures than those interacting with the flat bilayer.

In addition to differences in overall *β*-content, the spatial distribution of these structural features along the peptide sequence is markedly altered by curvature (Figure SI.10). In the flat bilayer system, *β*-structures are primarily localized within the central hydrophobic core (CHC). In contrast, for peptides bound to the vesicle, *β*-structures shift toward the other regions of the sequence. This redistribution likely arises because the CHC becomes engaged in strong interactions with the lipid hydrophobic core (Figure SI.3), reducing its availability for intramolecular hydrogen bonding. Notably, the metal-binding and polar regions – where a substantial fraction of *β*-structure is observed in the vesicle system – remain exposed to the solvent (Figure SI.3 (b)). As a result, although aggregation at the vesicle surface is suppressed, the peptides retain *β*-rich conformations that may induce conformational changes in incoming monomers via templating, as suggested in previous studies^13,22^.

An accounting of *β*-hairpin motifs, which are considered key intermediates in oligomer formation^68,69^, further highlights the changes in the A*β*42 conformational ensemble induced by membrane curvature. As shown in Figure 3 (b), *β*-hairpins in the flat bilayer system are only observed at early times, prior to adsorption, thus being associated with oligomerization in solution. In contrast, such motifs continue to appear after adsorption in the vesicle system. Not only are *β*-hairpins more likely to be found in the vesicle system compared to the flat bilayer, but the location of such motifs changes considerably. Residue-level analysis (Figure SI.11 (a) and (b)) reveals that, in the flat geometry, *β*-hairpins predominantly involve residues within the CHC and, to a lesser extent, the polar regions. For the vesicle, although *β*-hairpins primarily arise from the CHC region, there is more diversity in their locations, as other motifs also emerge in the metal-binding and C-terminal regions. This is in agreement with the observed redistribution of *β*-structure into these regions, as seen in Figure SI.10. All of these hairpin locations in the A*β*42 chain have been proposed in the literature, either from MD simulations or experimental data^55^. CHC – C-terminal hairpins, which have been traditionally implicated in oligomer formation ^55,68^, could not be detected, and may be hindered in the vesicle system due to a lack of contacts between these two regions after adsorption (see contact matrices in Figure SI.12). These are still retained to some extent in the flat bilayer simulations (Figure SI.13), although the CHC – C-terminal hairpin could also not be observed in this case. Nevertheless, the CHC – CHC and C-terminal – C-terminal hairpins have been shown to lead to oligomerization as well^70,71^. Importantly, *β*-hairpins within these regions remain accessible to the solvent (Figures SI.3 and SI.11 (c)), and may thus serve as nucleation sites for further peptide association with oncoming free A*β*42 peptides.

Additional evidence of curvature-induced structural differences in adsorbed A*β*42 is provided by the distribution of the radius of gyration (*R*_g_) (Figure SI.14). While both systems exhibit similar distributions prior to adsorption, the vesicle-bound peptides display a shift toward larger *R*_g_ values at later times, indicating more extended conformations. The only exception is A*β*42-2 in the flat bilayer system, which penetrates the membrane and remains monomeric rather than participating in trimer formation. This distinction underscores the difference between weakly adsorbed peptides that remain near the membrane surface and those that insert more deeply into the bilayer. Nevertheless, in spite of such similarity, the enhanced formation of *β*-structures and *β*-hairpin motifs is clearly associated with membrane curvature, as A*β*42-2 does not exhibit these features to a comparable extent (Figure 3).

### Properties of the lipid bilayers

In addition to modulating the adsorption and surface dynamics of A*β*42, membrane curvature also plays a key role in determining the structural response of the bilayer to peptide binding. Figure 4 illustrates the spatial distribution of lipids in the outer leaflet, which is directly exposed to A*β*42. In the vesicle system, a clear lipid raft emerges over the course of the simulation, characterized by the segregation of cholesterol and a large fraction of DPGS from the remaining lipid species. Cholesterol redistribution is further evidenced by the time evolution of the difference in cholesterol population between the inner and outer leaflets, as well as by the distance between the COM of cholesterol molecules and that of the entire vesicle (Figure SI.15 (b)). These results indicate that cholesterol not only forms a raft localized at one pole of the vesicle but also preferentially accumulates in the outer leaflet. On the other hand, the region surrounding the binding sites of A*β*42 is rich in PUPE and POPC lipids, while some DPGS still remains in contact with A*β*42, in line with the results shown in Figure SI.4.

**Figure 4.**
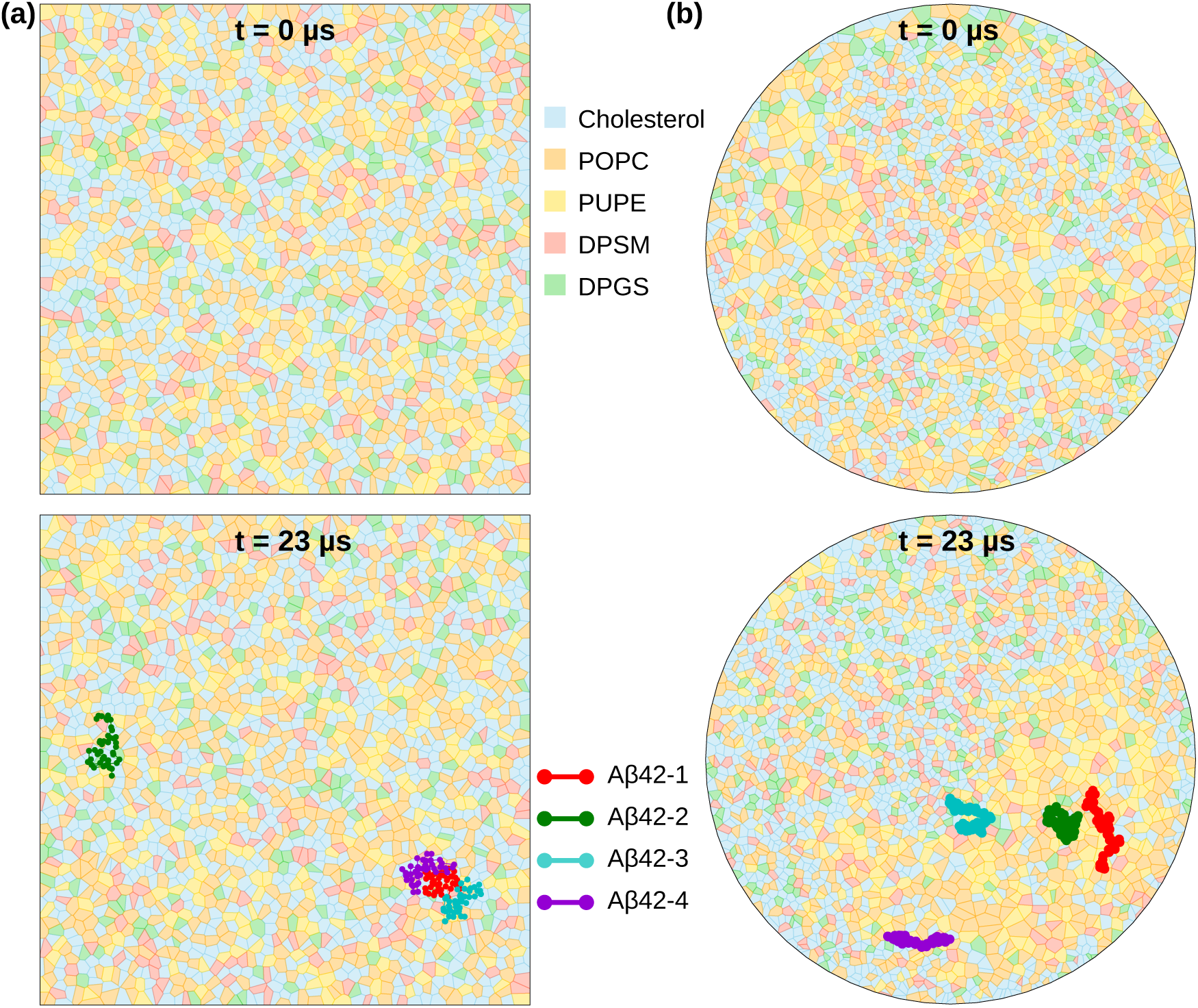
Distribution of lipids (Voronoi cells) and A*β*42 residues (colored points) on the outer leaflet of the membrane for the (a) flat bilayer system (top bilayer only) and (b) vesicle, at 0 and 23 µs. (a) represents normalized cartesian coordinates projected on the *xy* plane, whereas (b) represents polar coordinates in a Lambert azimuthal equal-area projection. Colors of each A*β*42 peptide match those in the trajectory frames shown in Figure SI.2.

It should be noted that intra-and interleaflet cholesterol migration are primarily driven by membrane curvature. Numerous studies have demonstrated a strong coupling between lipid raft dynamics and membrane curvature^72–75^. For instance, simulations by Pirhadi and Yong^76^ showed that leaflet asymmetry beyond a critical threshold induces stress that promotes the formation of segregated domains with distinct curvatures. Similarly, Gu et al.^77^ reported enhanced cholesterol flip-flop rates in less ordered membrane regions. No significant lipid segregation is observed in our simulatins of the flat bilayer system, where lipid distributions remain largely homogeneous (Figure 4a). Nonetheless, a slight enrichment of cholesterol in the upper leaflet occurs following A*β*42 adsorption (Figure SI.15a), along with a small divergence in APL between leaflets (Figure SI.6), suggesting that peptide binding may induce subtle, localized curvature changes.

Remarkably, in the vesicle system, the cholesterol-rich raft is located on the side opposite to the region where A*β*42 peptides are adsorbed (Figure 4 (b)). Furthermore, interleaflet cholesterol migration begins at approximately 1 µs, shortly after the initial peptide adsorption events (Figure 1 (b)), suggesting a correlation between peptide binding and lipid redistribution. It is difficult to conclusively determine to what extent A*β*42 directly drives raft formation compared to curvature, since some degree of cholesterol segregation is already present in the peptide-free vesicle (Figure SI.19 (c)). Nevertheless, there are three key observations that strongly indicate the ability of A*β*42 to modulate lipid organization on curved surfaces: (i) depletion of cholesterol from the peptide binding region, where it was initially present – see Figure SI.19 (b) and also the initial bump in the profile of A*β*42 – cholesterol interactions in Figure 1 (b); (ii) spatial exclusion of the raft from the peptide-bound region at equilibrium and (iii) the observed interleaflet migration of cholesterol after peptide adsorption.

Figures 1 and SI.4 suggest a possible mechanism for this behavior. A*β*42 interacts more favorably with DPGS, PUPE and POPC than with cholesterol – both via electrostatic and hydrophobic interactions –, effectively recruiting these lipids to the peptide-bound region. These interactions also induce local membrane deformation, as lipids near A*β*42 were found to be displaced outward relative to the average headgroup position of the outer leaflet (Figure SI.16), whereas the opposite occurred in the raft. The formation and peptide-induced reorganization of the raft is also facilitated by the intrinsically higher disorder of the vesicle membrane, reflected in the lower *S*_C_*_D_* values of lipid acyl chains compared to the flat bilayer (Figure SI.17). The increased disorder enhances membrane fluidity, enabling large-scale lipid rearrangements.

Lipid rafts are typically liquid-ordered (L_o_) domains, coexisting with liquid-disordered (L_d_) regions, and are characterized by increased membrane thickness^73,78^. Our results are consistent with this picture: the raft exhibits higher local *S*_C_*_D_* values and greater thickness (Figure SI.18), identifying it as a cholesterol-rich L_o_ phase, while the surrounding domain corresponds to L_d_ phase, rich in polyunsaturated PUPE. As such, A*β*42 displays a clear preference for the L_d_ region of the curved membrane. In fact, such an effect has been demonstrated experimentally by Staneva et al. ^79^ through fluorescence studies using giant and large unilamellar vesicles (LUVs), by Ahyayauch et al. ^80^ using calorimetric measurements with both LUVs and flat bilayers, and Azouz et al.^81^ via atomic force microscopy (AFM). Interestingly, there is evidence in the work of Staneva et al. that A*β*42 binding to the L_d_ domains of the membrane resulted in coalescence of L_o_ domains^79^. This aligns with our findings showing A*β*42 and L_o_ anti-localization in the vesicle. Even the AFM imaging studies of Azouz et al. have noted a slight homogeneous coalescence of the L_o_ and L_d_ of a flat bilayer in the presence of A*β*42.

At first glance these findings may appear to contradict reports of A*β*42 affinity for lipid rafts, attributed to favorable interactions with cholesterol and GM1 gangliosides^82–84^. However, as pointed out by Staneva et al., preference for L_d_ domains does not preclude binding to L_o_ phases^79^. Conversely, although the lipid rafts may serve as regions where A*β*42 production is enhanced, due to higher activity of *β* and *γ*-secretases^83,84^, this does not necessarily imply that monomeric A*β*42 must be confined to them^80^. Partition of A*β*42 between the domains is likely governed by a balance between A*β*42 interactions with cholesterol and glycosphingolipids and hydrophobic interactions with lipid acyl chains. In curved membranes, the increased exposure of acyl chains due to packing defects favors the latter, dominating peptide binding behavior—a mechanism absent in flat bilayers.

Furthermore, three key points are worth addressing. First, it must be reiterated that the DPGS glycosphingolipid did not fully segregate into the raft domain, and was still available in the L_d_ where it interacted strongly with A*β*42. Staneva et al. also showed that presence of GM1 on the L_d_ phase modulates its response to A*β*42 binding, which leads to greater penetration and increase in disorder, when compared to a L_d_ phase without GM1^79^. Second, cholesterol molecules were not completely absent from the process of A*β*42 binding. In Figure 1 (b), the initial increase in A*β*42 – cholesterol hydrophobic contacts between 0 and 4 µs suggests that cholesterol initially present in the binding region helped recruit A*β*42 to the surface, before segregating into the raft. Third, it has been suggested that the boundary between the L_o_ and L_d_ may serve as key features for A*β*42 binding to the membrane^81,85,86^. Indeed, in Figure 4 (b) it can be seen that two of the A*β*42 peptides (3 and 4) are located precisely at the interface of the two domains.

The link between A*β*42 adsorption and membrane packing defects is further highlighted by the spatial distribution of local APL values (Figure SI.19). In the vesicle, regions surrounding the adsorbed peptides exhibit significantly higher APL compared to those of the lipid raft. Nevertheless, the packing defects have merely relocated on the surface, as their area distribution (and total area) did not change significantly compared to a vesicle without adsorbed A*β*42 – see Figures SI.7 and 2.

Interestingly, a similar (albeit more localized) effect is observed in the flat bilayer system: a small region of increased APL coincides with the adsorption site of A*β*42-2, the only peptide that penetrates the membrane, as seen in Figure SI.19 (c). Furthermore, the planar bilayer system exhibits a distribution of packing defect areas slightly shifted to higher values. This effect is likely associated with the aforementioned insertion of A*β*42-2 and the changes in APL of the flat membrane. It suggests that A*β*42 can induce local disruptions even in planar membranes, enlarging packing defects and thus promoting access to the hydrophobic core, which in turn facilitates deeper insertion. The enlargement of packing defects in planar supported bilayers by A*β*42 has in fact been recently observed by AFM^87^.

## Conclusion

Through all-atom molecular dynamics simulations, we have elucidated the role of lipid membrane curvature in modulating A*β*42 dynamics and its interplay with lipid bilayers. In planar bilayers, A*β*42 interactions are relatively weak and largely confined to lipid headgroups. Adsorbed peptides remain near the membrane surface, where dimers can diffuse laterally and grow into larger oligomers, such as trimers. By contrast, on curved vesicle membranes, A*β*42 monomers penetrate more deeply by accessing the hydrophobic core through packing defects. This enables strong interactions with lipid acyl chains, primarily via the CHC and C-terminal regions. These interactions stabilize A*β*42 in more extended conformations enriched in aggregation-prone *β*-structures, including *β*-hairpins, to a greater extent than observed on planar membranes. Importantly, these structural motifs are located in regions of the peptide that remain solvent-exposed, suggesting a potential ability to template conformational changes in incoming monomers. While this mechanism was not directly demonstrated here, it represents an important direction that motivates future investigation.

Our results further reveal that membrane curvature governs its structural response to peptide adsorption. In the vesicle system, the high curvature promotes the emergence of a cholesterol-rich liquid-ordered (L_o_) domain, while A*β*42 preferentially localizes to the opposing liquid-disordered (L_d_) region enriched in phosphoethanolamine lipids. Peptide adsorption is accompanied by net cholesterol migration toward the outer leaflet, highlighting a strong coupling between peptide binding and lipid organization. On the other hand, the planar bilayer does not exhibit large-scale domain formation, although localized expansion of packing defects is observed. By preferentially binding to L_d_ regions on curved membranes, A*β*42 may achieve higher local surface concentrations compared to planar systems, with potential enhancement of on-surface oligomerization processes.

Several aspects of curvature remain to be explored, including the effects of curvature magnitude and sign (concave versus convex geometries). Nonetheless, this work demonstrates that membrane curvature is a critical factor in surface-catalyzed A*β*42 aggregation and should not be neglected. By promoting the formation of hydrophobic packing defects that act as nucleation sites for misfolded conformations, highly curved membranes may play a key role in the early stages of toxic oligomer formation. These findings suggest that membrane physical properties, in addition to molecular composition, should be considered when identifying potential therapeutic targets aimed at modulating A*β*42 oligomerization.

## Declaration on the use of generative AI and AI-assisted technologies

The authors used ChatGPT exclusively to assist with Python code development, involved in trajectory analysis, and for language editing, to improve the manuscript’s clarity and readability. All code outputs were verified and validated by the authors. No generative AI tools were used for the data analysis, interpretation, or drafting of the scientific content. The authors are fully responsible for all the content and conclusions presented.

## Acknowledgement

Anton 3 computer time was provided by the Pittsburgh Supercomputing Center (PSC) through Grant 1R24GM154042 from the National Institutes of Health. The Anton 3 machine at PSC is made available by D. E. Shaw Research. The authors would also like to thank Filipe Carvalho for helpful contributions to some of the analysis and parts of the text.

## Supporting Information Available

Additional figures and tables.

