## Supplementary material for "Ahead of the membrane curve: *in silico* insights into amyloid-*β* aggregation": SI

### Supporting Information

Pedro Maximiano and Mohtadin Hashemi*

*Department of Physics, Auburn University, Leach Science Center, Auburn, Alabama 36849*

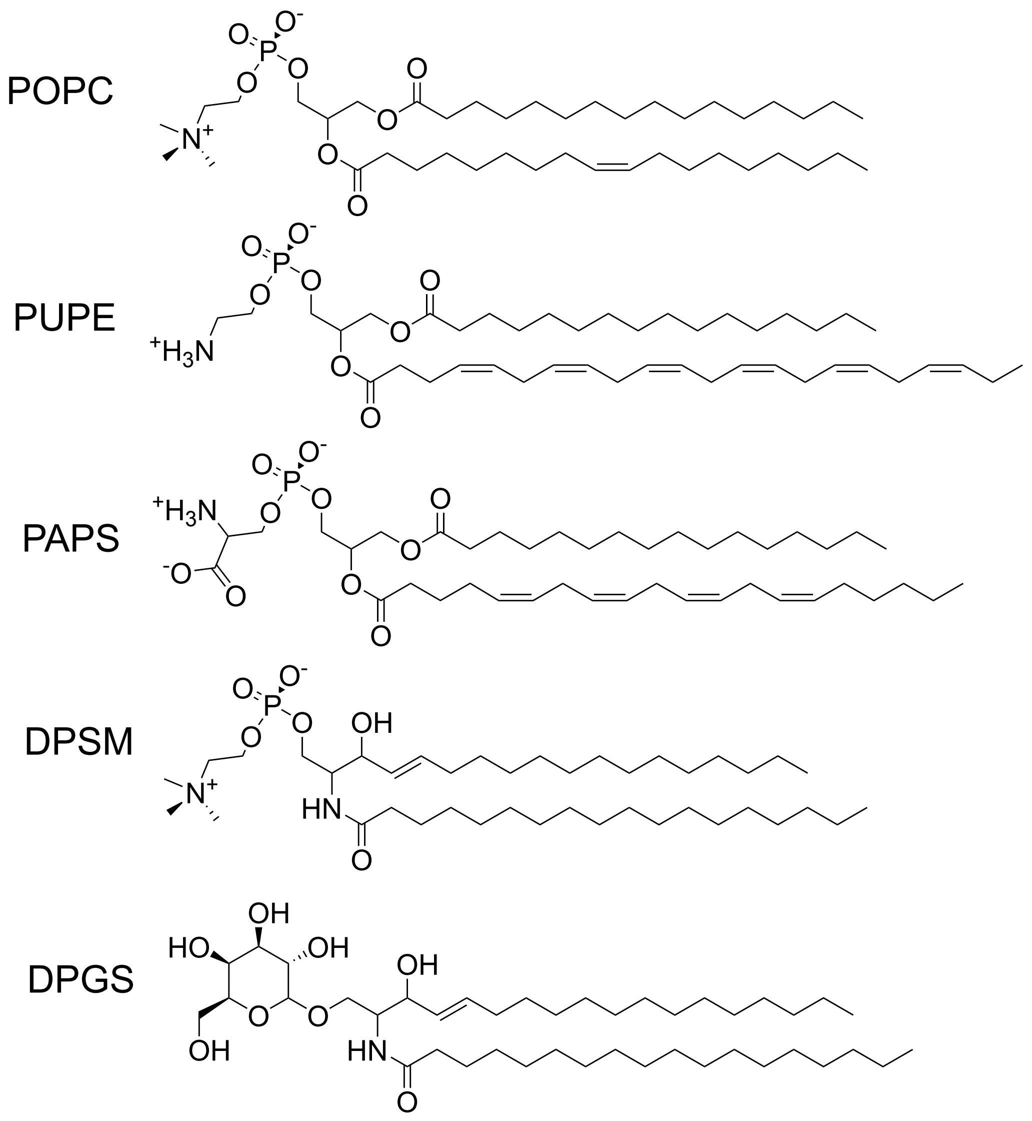


**Figure SI.1.** Structure of the phospho- and sphingolipid species that make up the bilayer models employed in this work.

**Table SI.1:** Molecular composition of the simulated systems.

| Species | | | Number of molecules | | | | |
|---|---|---|---|---|---|---|---|
| | | | Flat bilayer | Flat bilayer + Aβ42 | Vesicle, initial | Vesicle, equilibrated | Vesicle + Aβ42 |
| Lipid bilayer | Outer leaflet | Cholesterol | 1786 | 1786 | 1009 | 1036 | 1039 |
| | | POPC | 842 | 842 | 593 | 593 | 593 |
| | | PUPE | 736 | 736 | 265 | 265 | 265 |
| | | PAPS | 234 | 234 | 0 | 0 | 0 |
| | | DPSM | 244 | 244 | 215 | 215 | 215 |
| | | DPGS | 160 | 160 | 180 | 180 | 180 |
| | Inner leaflet | Cholesterol | 1786 | 1786 | 691 | 664 | 661 |
| | | POPC | 842 | 842 | 246 | 246 | 246 |
| | | PUPE | 736 | 736 | 389 | 389 | 389 |
| | | PAPS | 234 | 234 | 181 | 181 | 181 |
| | | DPSM | 244 | 244 | 42 | 42 | 42 |
| | | DPGS | 160 | 160 | 0 | 0 | 0 |
| Intracellular domain | | Water | 322452 | 322452 | 52209 | 52209 | 52183 |
| | | Na+ | 528 | 528 | 92 | 92 | 92 |
| | | K+ | 4531 | 4531 | 799 | 799 | 799 |
| | | Glutmate | 4591 | 4591 | 710 | 710 | 710 |
| Extracellular domain | | Water | 353119 | 337198 | 1001090 | 1001090 | 948907 |
| | | Na+ | 5280 | 5305 | 920 | 920 | 932 |
| | | K+ | 151 | 152 | 27 | 27 | 29 |
| | | Cl- | 5431 | 5444 | 947 | 947 | 948 |
| | | Aβ42 | 0 | 4 | 0 | 0 | 4 |

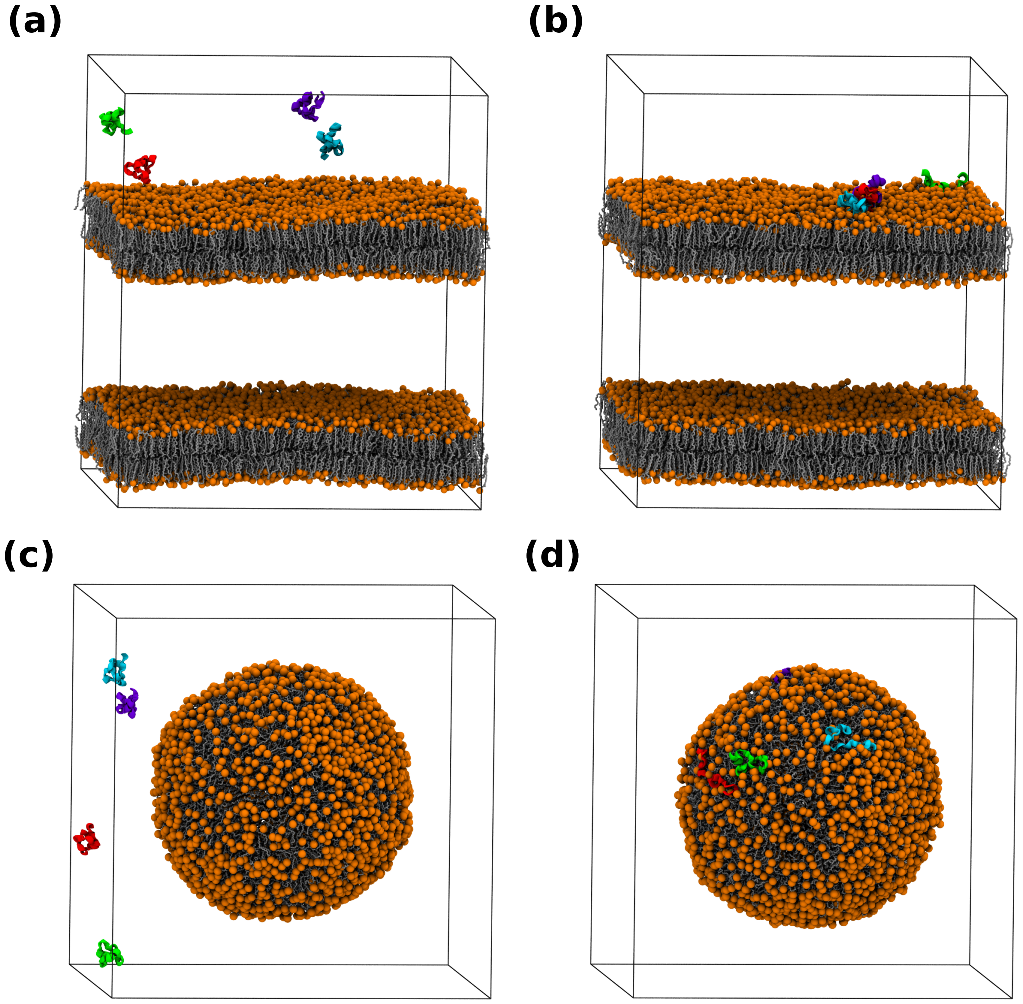


**Figure SI.2.** Frames extracted from the flat bilayer system, at $t = 0.0\,\mu s$ (a) and $t = 23.0\,\mu s$ (b), and from the vesicle system at $t = 0.0\,\mu s$ (c) and $t = 23.0\,\mu s$ (d). Water and ions omitted for clarity.

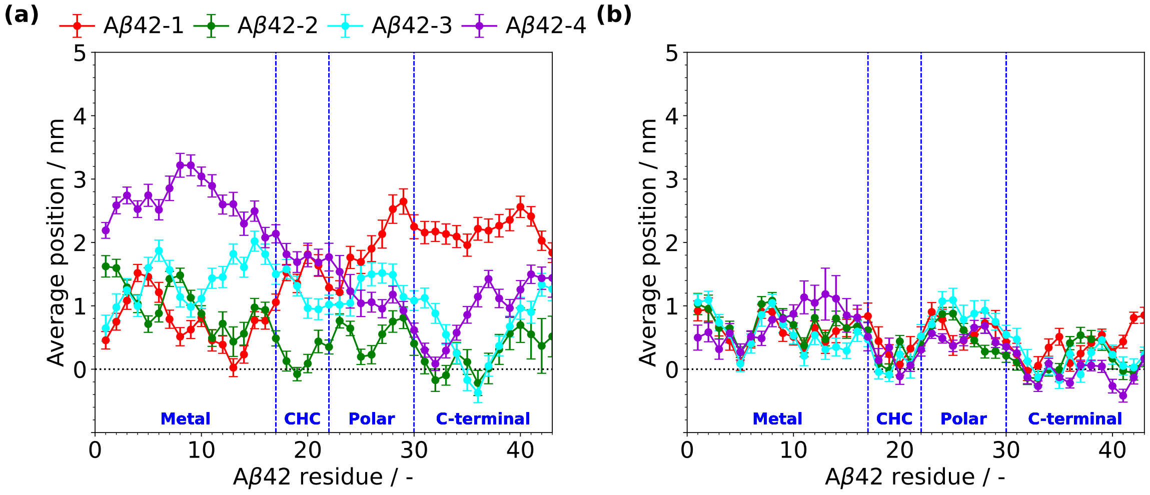


**Figure SI.3.** Average position of the center-of-mass (COM) of each Aβ42 residue in the direction transversal to the lipid bilayer, relative to the average position of the outer leaflet headgroup atoms (dotted black line), in the flat bilayer (a) and vesicle systems (b). The averages were calculated for the last 1.0 µs of trajectory, from samples taken every 100 ns to avoid time correlation. Colors of each Aβ42 peptide match those in the trajectory frames shown in Figure SI.2.

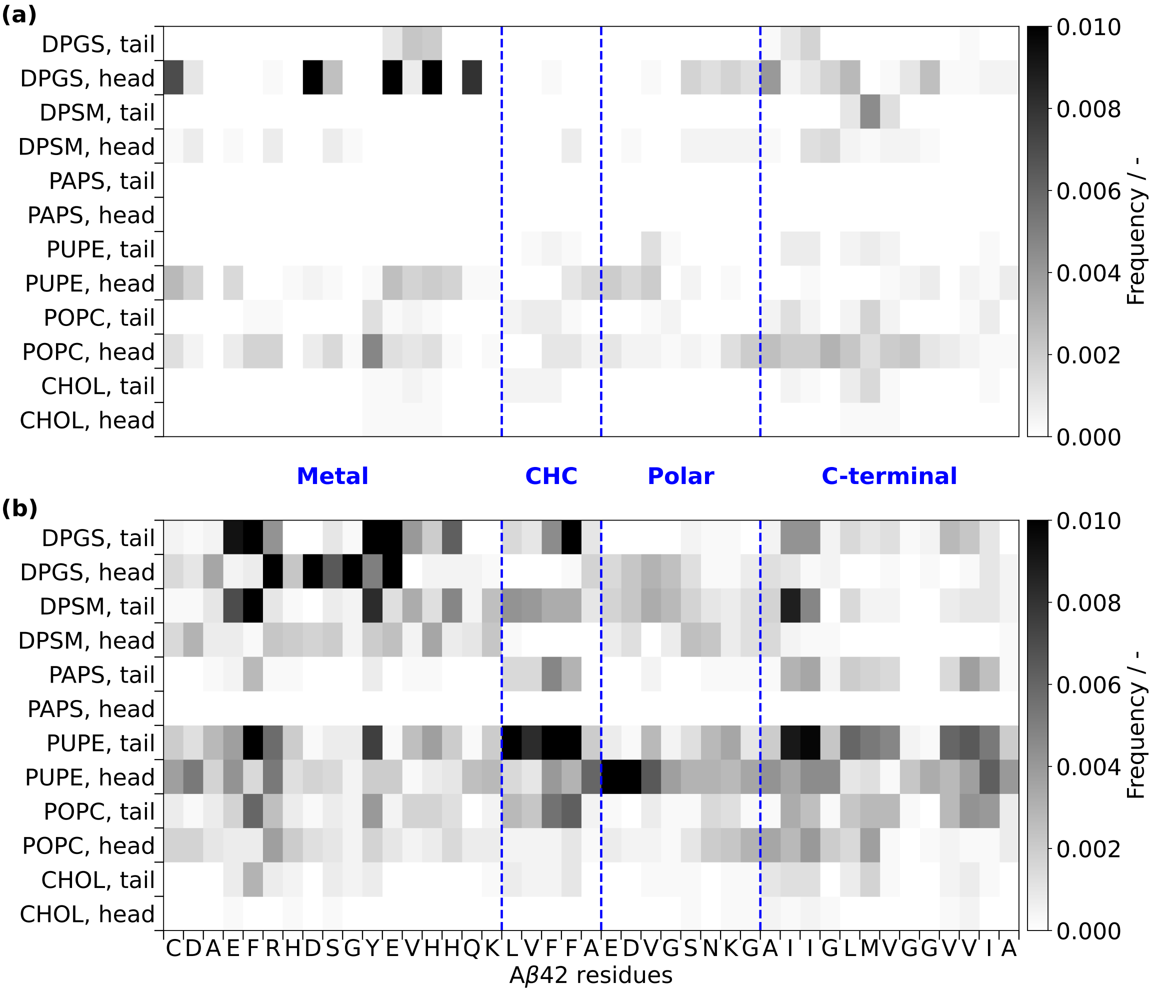


**Figure SI.4.** Lipid–Aβ42 contact matrices for the flat bilayer (a) and vesicle systems (b), calculated for the last 1.0 µs of trajectory.

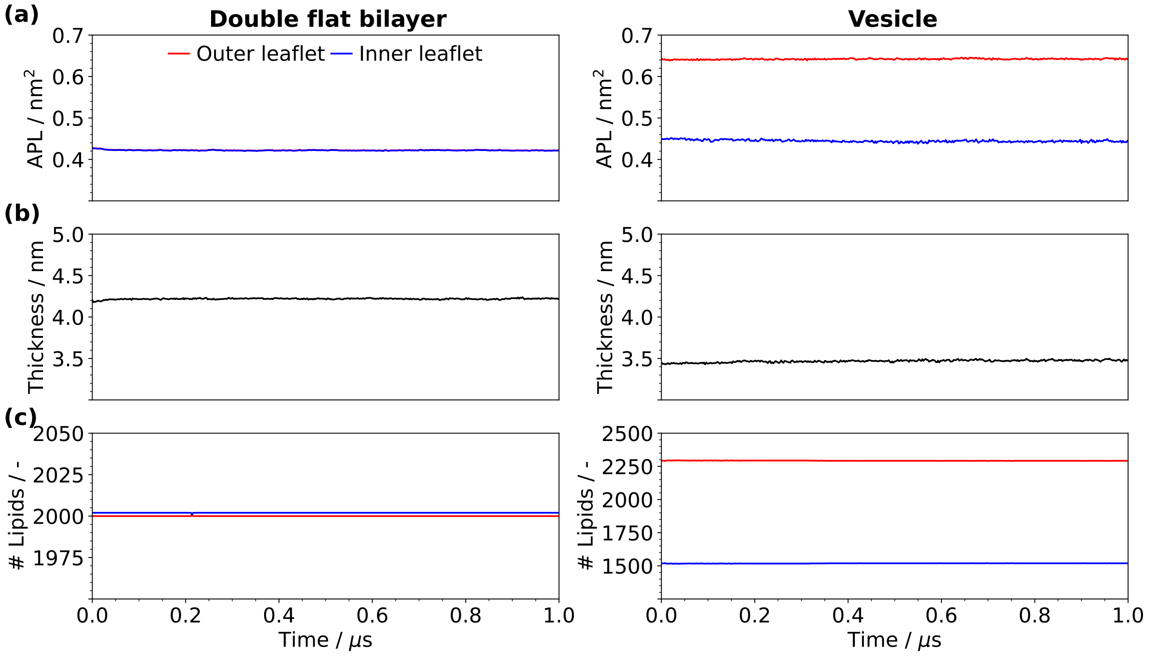


**Figure SI.5.** Area per lipid (APL) (a), thickness (b) and number of lipid molecules (c) of the top flat bilayer (left) and the vesicle (right) systems simulated without Aβ42.

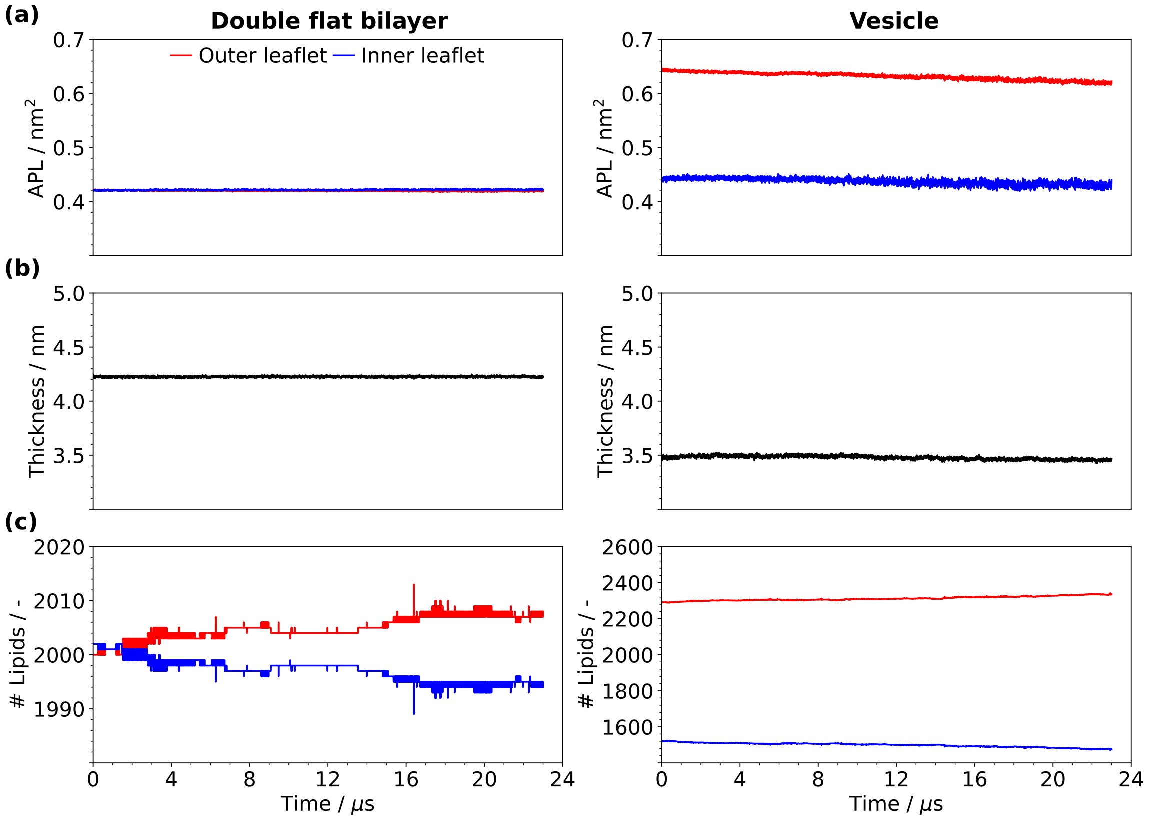


**Figure SI.6.** APL (a), thickness (b) and number of lipid molecules (c) of the top flat bilayer (left) and the vesicle (right), for the systems containing Aβ42.

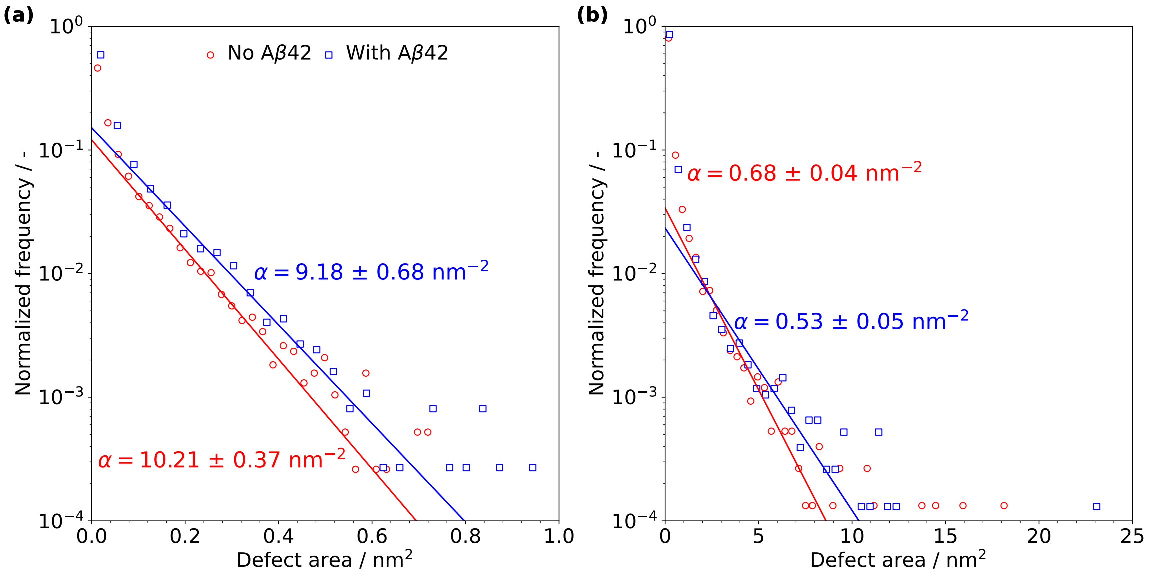


**Figure SI.7.** Distribution of hydrophobic packing defect areas for the flat bilayer (a) and vesicle (b) systems. The points correspond to binned data in an histogram, to which an exponential distribution of the form $p(x) = C\exp(-\alpha x)$ was fitted.

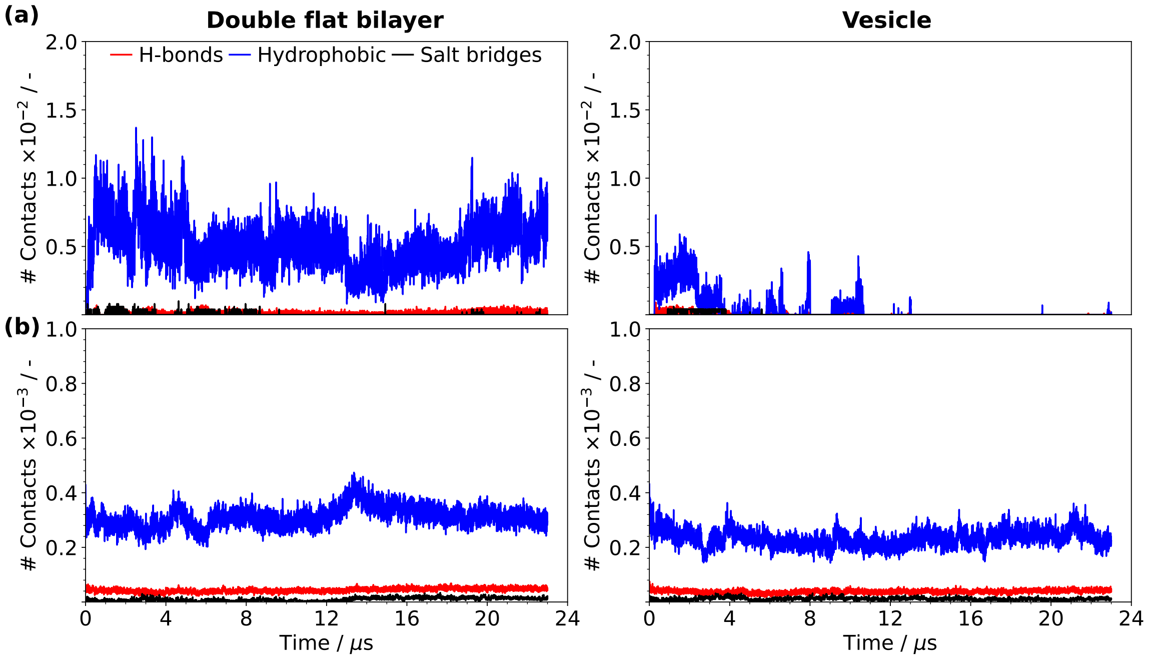


**Figure SI.8.** Number of intermolecular (a) and intramolecular (b) interactions – H-bonds, hydrophobic contacts and salt bridges – involving Aβ42 peptides as a function of simulation time for the flat bilayer (left) and vesicle (right) trajectories.

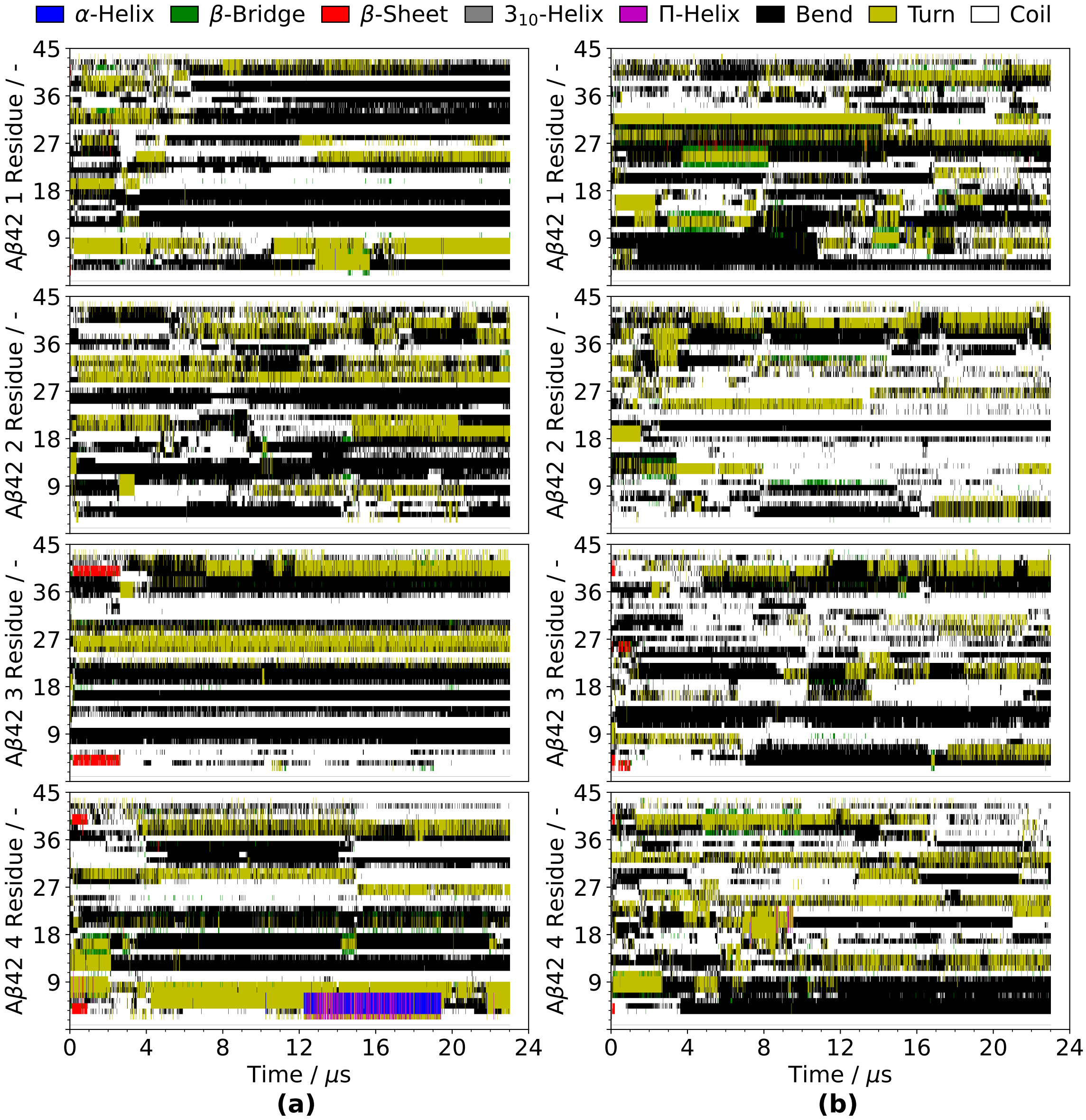


**Figure SI.9.** Secondary structure of Aβ42 residues as a function of simulation time for the flat bilayer (a) and vesicle (b) systems.

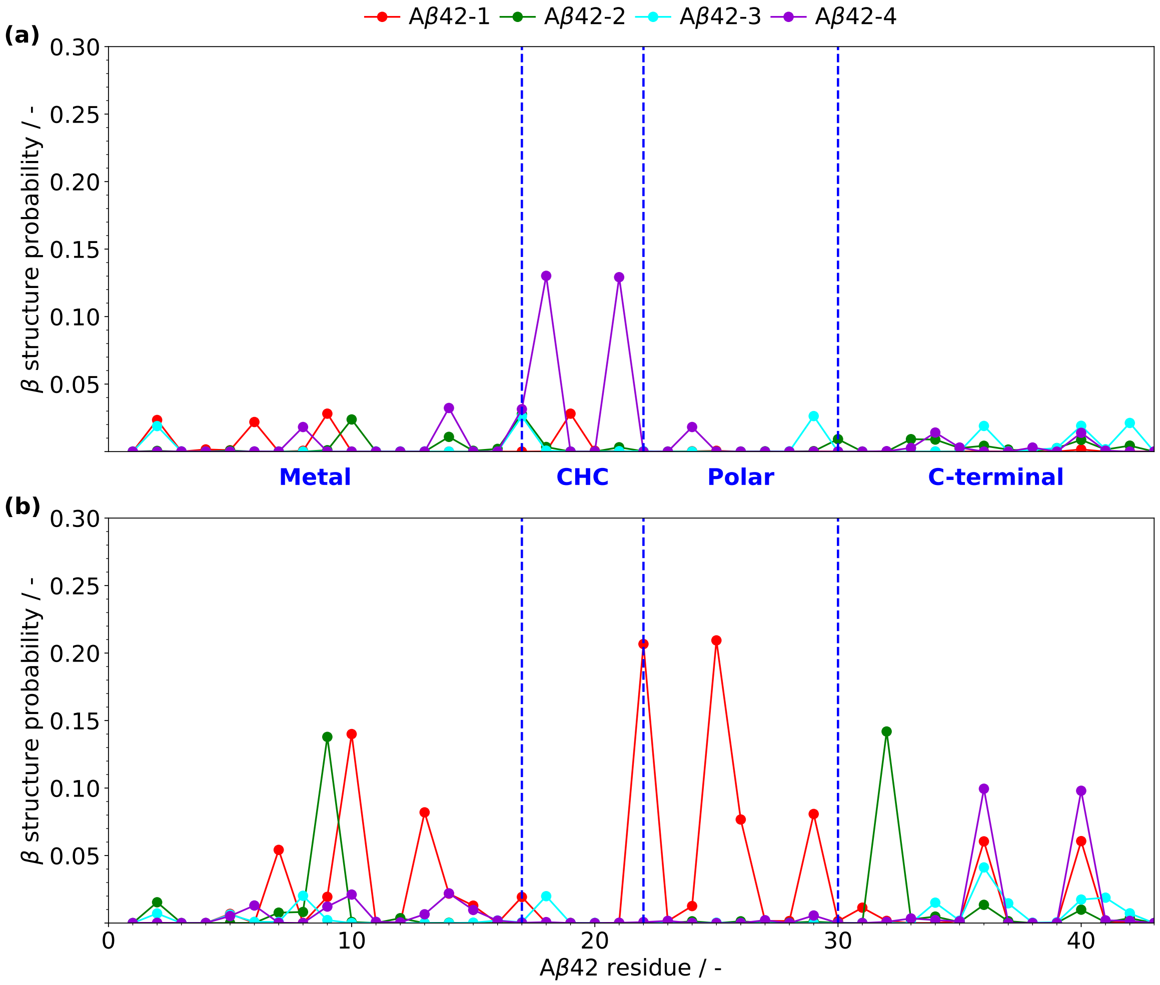


**Figure SI.10.** Probability of Aβ42 residue membership in a β structure of any kind for the flat bilayer (a) and vesicle (b) systems. Data was accumulated only between 4.0 and 23.0 μs, to reflect the structure of adsorbed Aβ42 peptides.

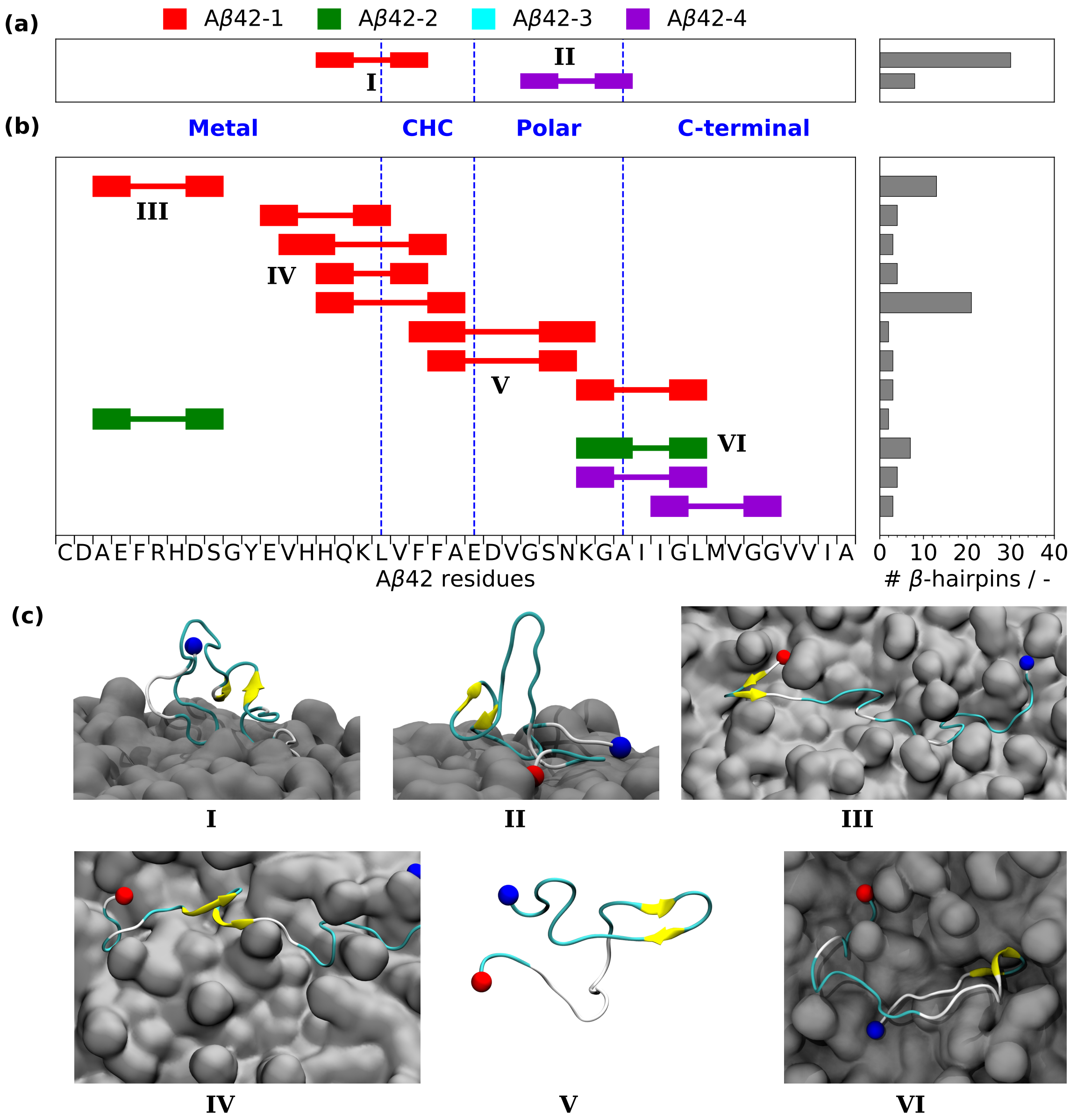


**Figure SI.11.** Per-residue accounting of β hairpin motifs detected in each Aβ42 peptide on the flat bilayer (a) and vesicle (b) trajectories (total number of motifs for the entire trajectory). (c) Structures of the most prevalent β hairpins detected – roman numeral labels correspond to different hairpin groups on (a) and (b). Red and blue spheres represent the N and C termini of Aβ42, respectively.

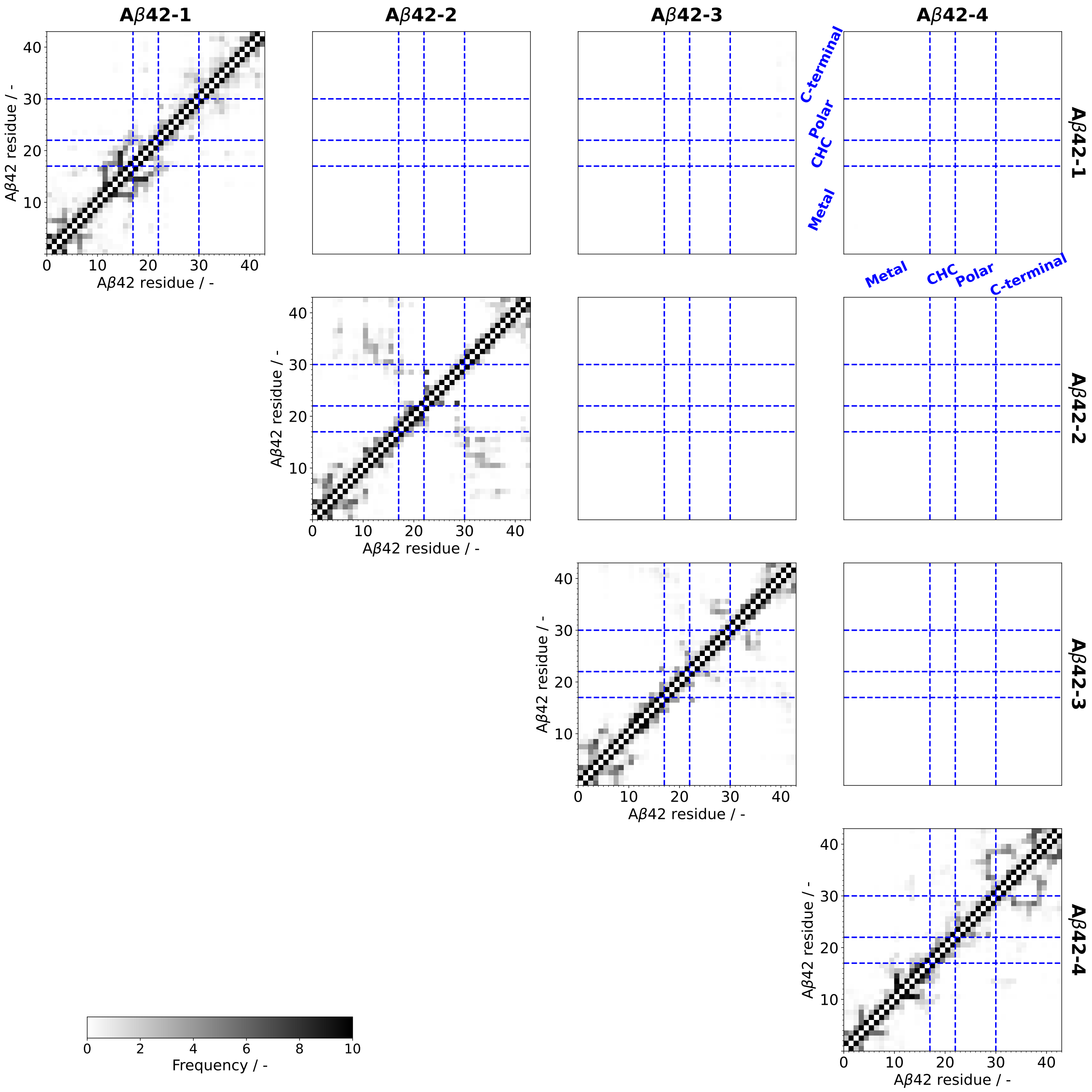


**Figure SI.12.** Aβ42–Aβ42 contact matrices for the interval 4.0 – 23.0 µs of the vesicle trajectory.

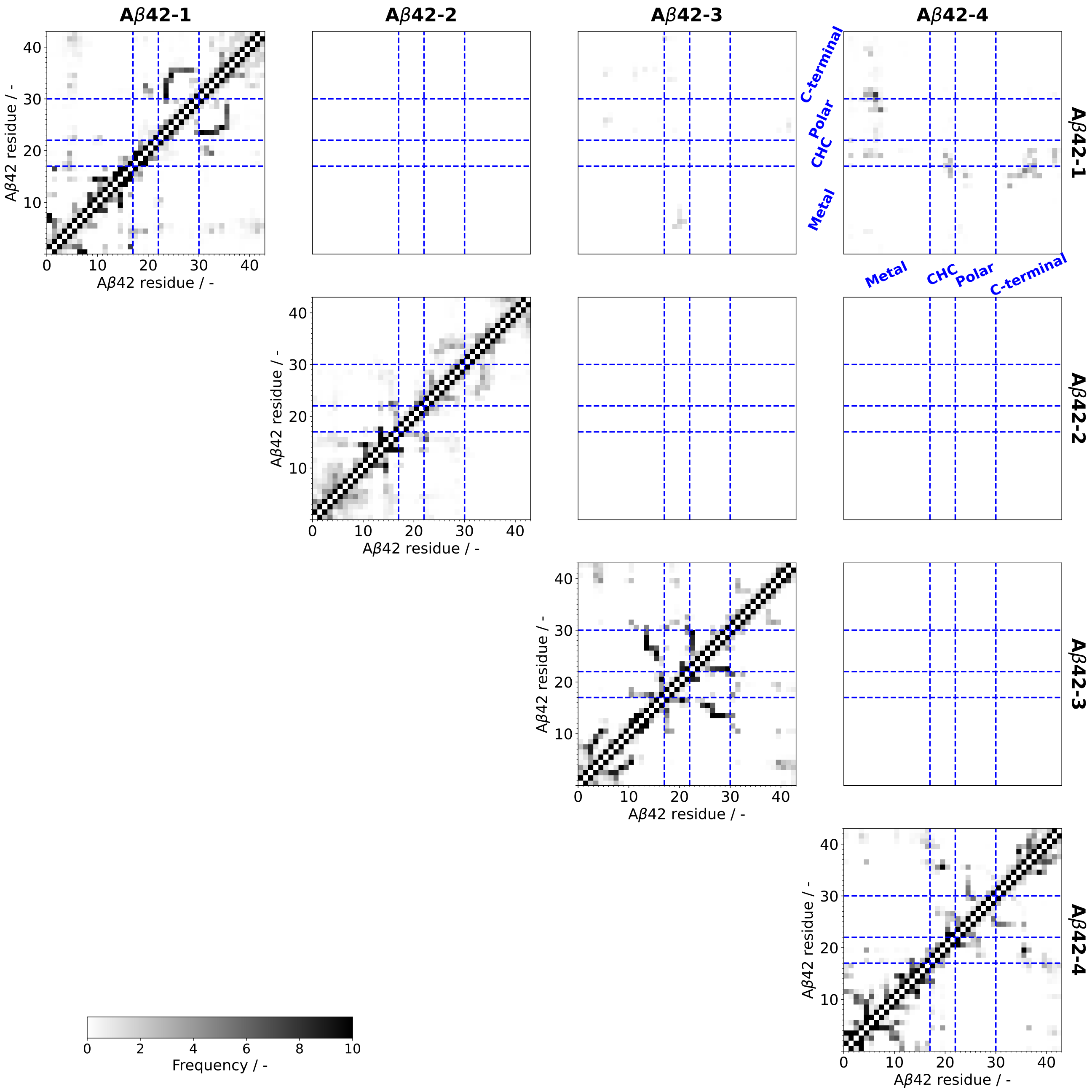


**Figure SI.13.** Aβ42–Aβ42 contact matrices for the interval 4.0 – 23.0 µs of the flat bilayer trajectory.

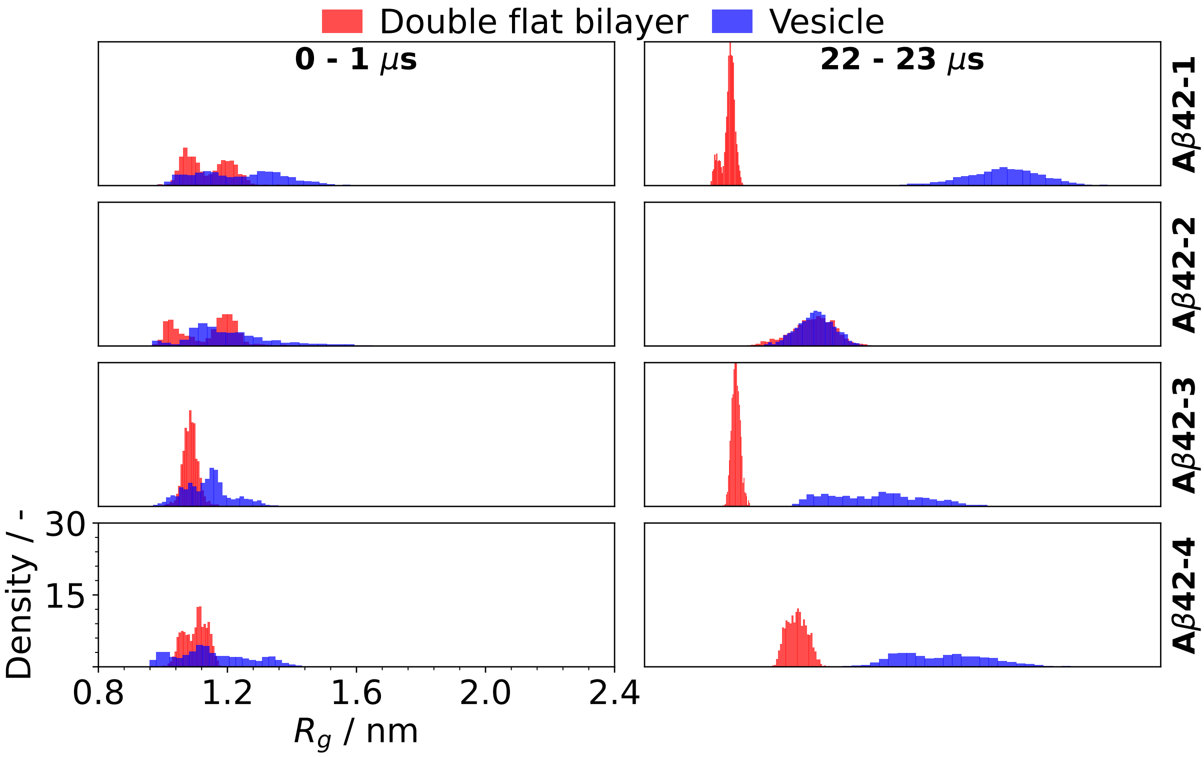


**Figure SI.14.** Distributions of the values of radius of gyration ($R_\mathrm{g}$) of each Aβ42 peptide for the first and last 1.0 us of the trajectories of each system.

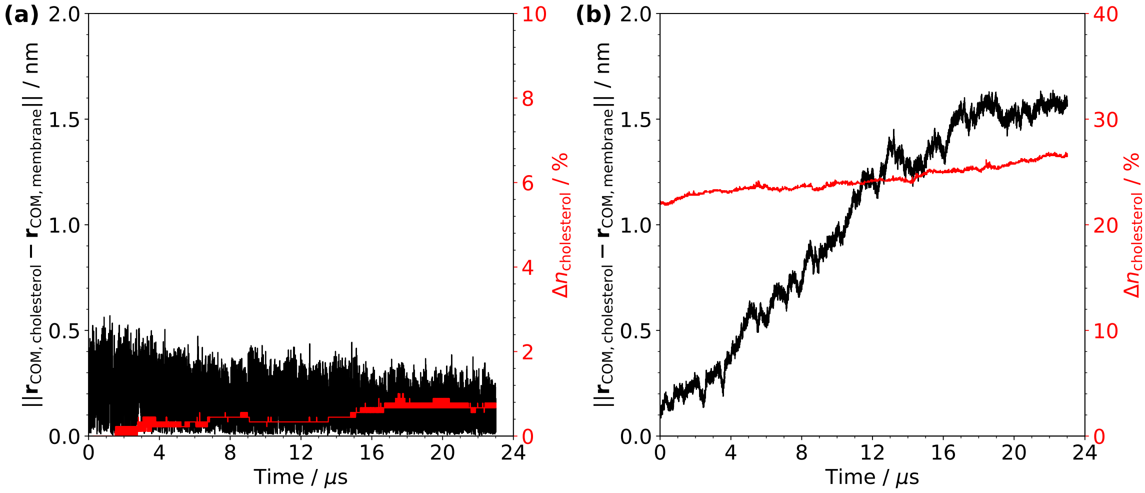


**Figure SI.15.** Distance between the COM of cholesterol lipids and the COM of the bilayer (black) and relative leaflet cholesterol number mismatch ($\Delta n_{\mathrm{cholesterol}}$, red) for the flat bilayer (a) and vesicle (b) systems. $\Delta n_{\mathrm{cholesterol}} = 100 \times (n_\mathrm{o} - n_\mathrm{i})/(n_\mathrm{o} + n_\mathrm{i})$, where $n_\mathrm{o}$ and $n_\mathrm{i}$ are the number of cholesterol molecules in the outer and inner leaflets, respectively.

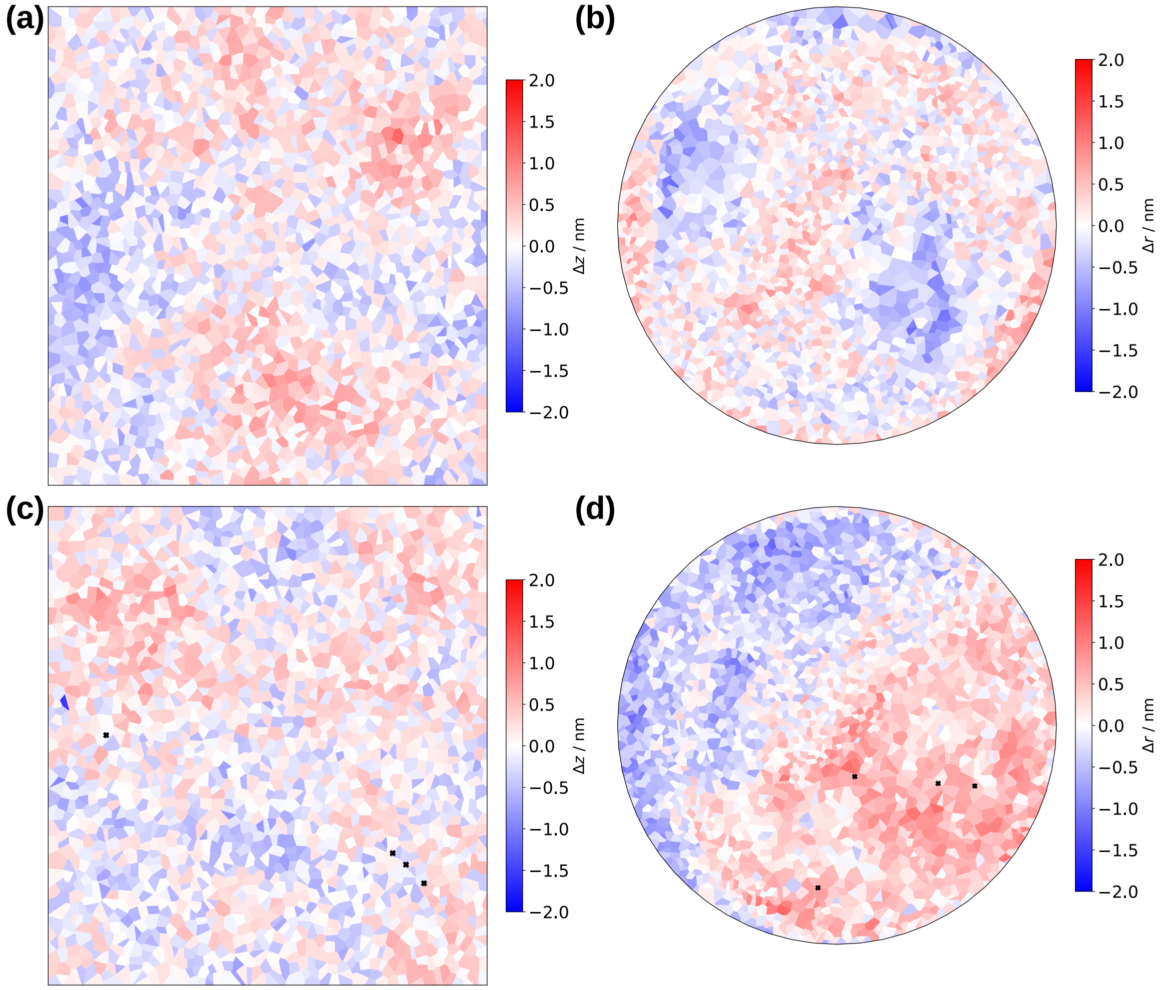


**Figure SI.16.** Local transversal deviation of lipid head group positions from the average of the outer leaflet on the top flat bilayer – (a) and (c) – and the vesicle – (b) and (d) – at the last frame. (a) and (b) refer to systems without Aβ42 and (c) and (d) are systems with Aβ42 (crosses mark the positions of Aβ42 COMs). (a) and (c) represent normalized cartesian coordinates projected on the $xy$ plane, whereas (b) and (d) represent polar coordinates in a Lambert azimuthal equal-area projection.

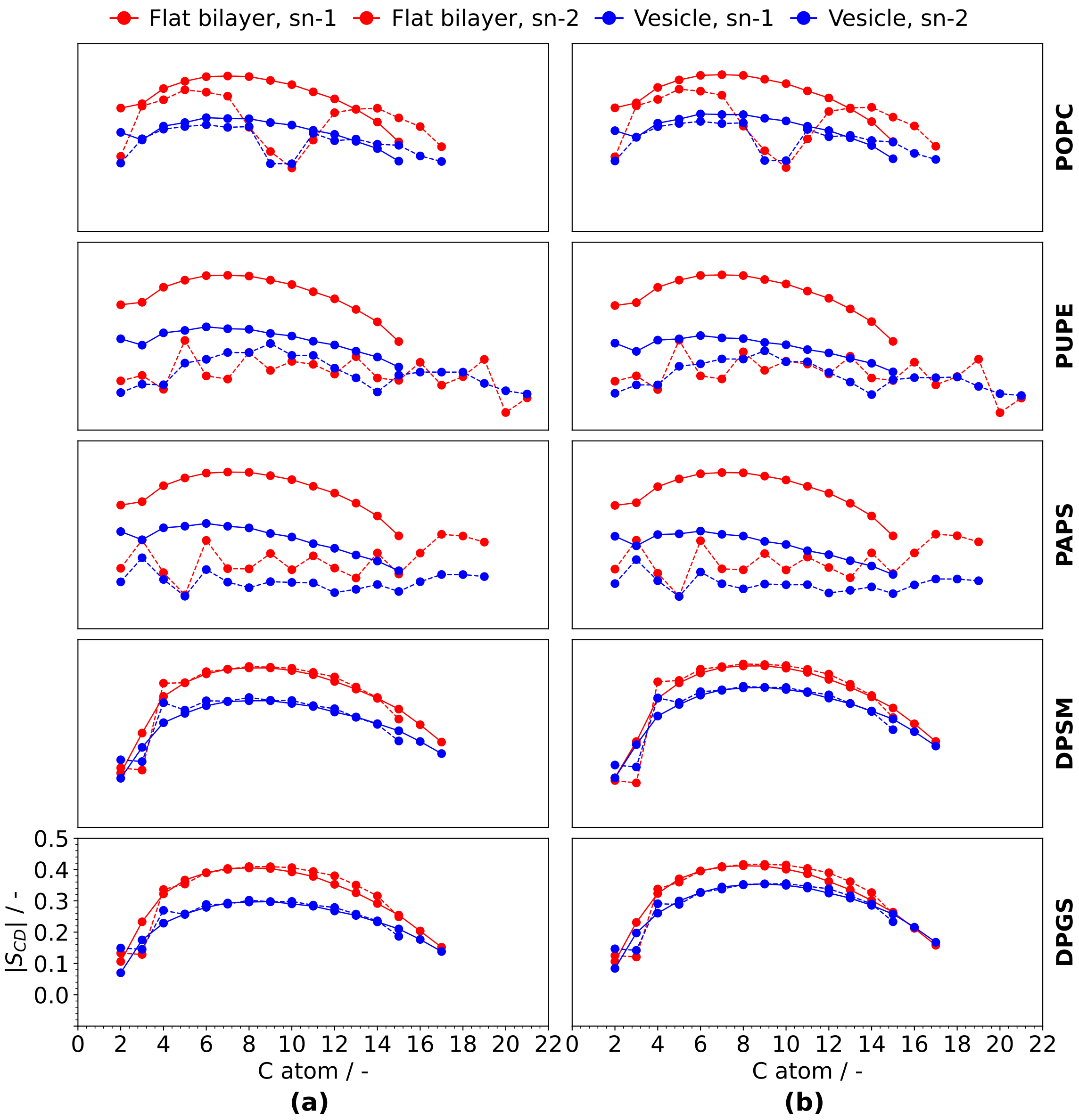


**Figure SI.17.** Average lipid order parameters, calculated for the last 1 µs of trajectory for the systems without Aβ42 (a) and with Aβ42 (b).

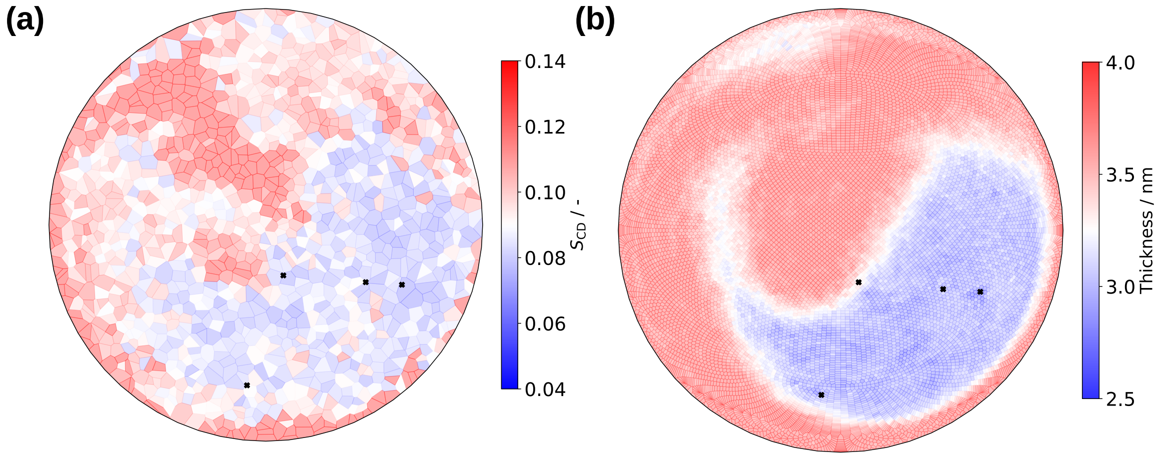


**Figure SI.18.** Local average lipid order parameters (a) and thickness (b), calculated for the last 1 µs of trajectory for the vesicle system with Aβ42. In (a), $S_{\mathrm{CD}}$ values were calculated via Equation 1 by averaging over multiple frames and all tail C atoms of each lipid, and are represented on a Voronoi tesselation done on the final lipid head group positions, in a Lambert azimuthal equal area projection. In (b), local thickness values were calculated by projecting lipid head group positions on a HEALPix grid drawn on the vesicle membrane midplane (resolution of 16 Å$^2$). An average thickness was then calculated on each grid cell from the radii of the inner and outer leaflet lipids belonging to it.

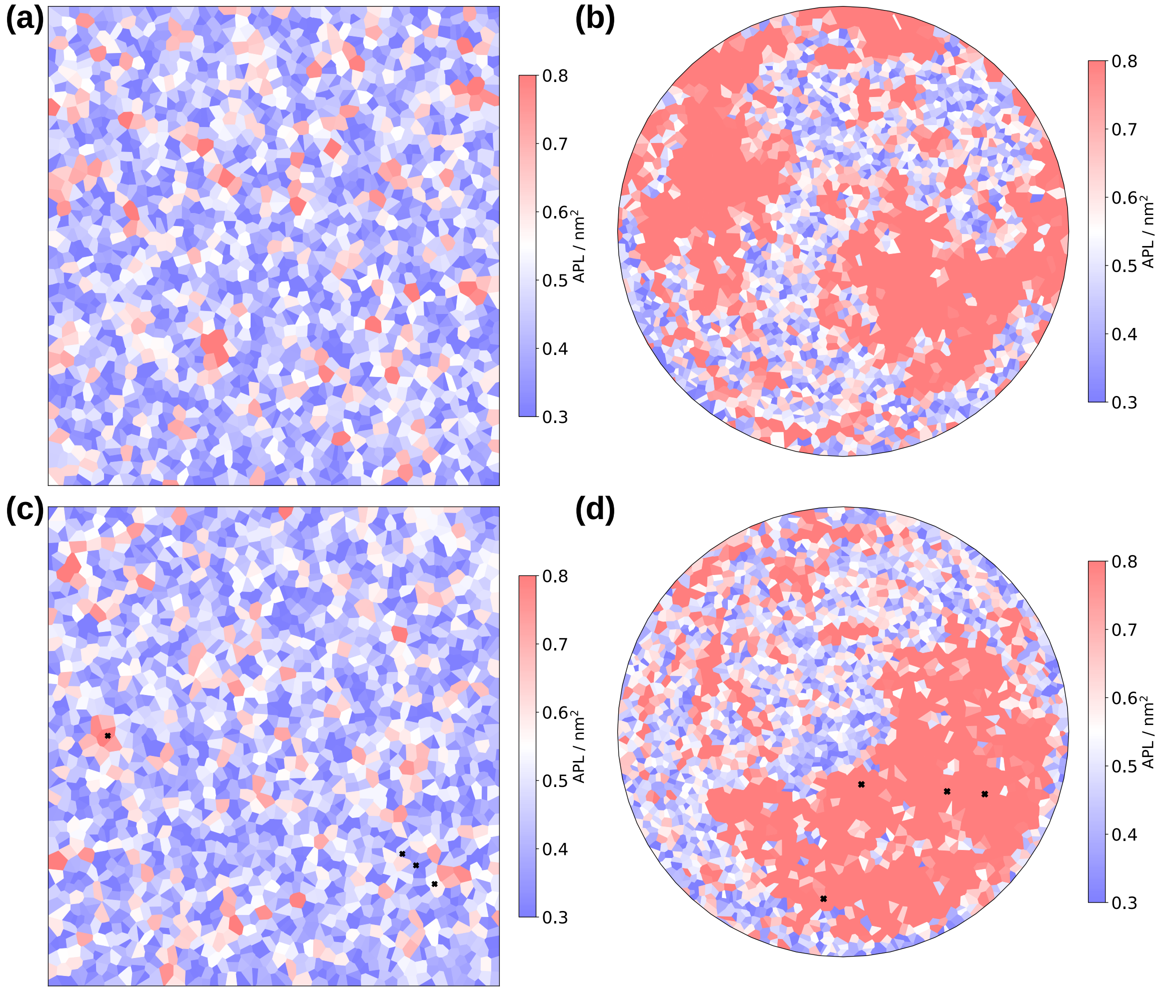


**Figure SI.19.** Local APL distribution on the outer leaflet of the top flat bilayer – (a) and (c) – and the vesicle – (b) and (d) – at the last frame. (a) and (b) refer to systems without Aβ42 and (c) and (d) are systems with Aβ42 (crosses mark the positions of Aβ42 COMs). (a) and (c) represent normalized cartesian coordinates projected on the $xy$ plane, whereas (b) and (d) represent polar coordinates in a Lambert azimuthal equal-area projection.
